# Luminosity thresholds of colored surfaces are determined by their upper-limit luminances empirically internalized in the visual system

**DOI:** 10.1101/2021.06.21.449225

**Authors:** Takuma Morimoto, Ai Numata, Kazuho Fukuda, Keiji Uchikawa

## Abstract

We typically have a fairly good idea whether a given object is self-luminous or illuminated, but it is not fully understood how we make this judgement. This study aimed to identify determinants of the luminosity threshold, a luminance level at which a surface begins to appear self-luminous. We specifically tested a hypothesis that our visual system knows the maximum luminance level that a surface can reach under the physical constraint that a surface cannot reflect more light than any incident light and applies this prior to determine the luminosity thresholds. Observers were presented with a 2-degree circular test field surrounded by numerous overlapping colored circles and luminosity thresholds were measured as a function of (i) the chromaticity of the test field, (ii) the shape of surrounding color distribution and (iii) the color of the illuminant of the surrounding colors. We found that the luminosity thresholds peaked around the chromaticity of test illuminants and decreased as the purity of the test chromaticity increased. However, the loci of luminosity thresholds across chromaticities were nearly invariant to the shape of the surrounding color distribution and generally resembled the loci drawn from theoretical upper-limit luminances and upper-limit luminance boundaries of real objects. These trends were particularly evident for illuminants on the black-body locus and did not hold well under atypical illuminants such as magenta or green. These results support the idea that our visual system empirically internalizes the gamut of surface colors under natural illuminants and a given object appears self-luminous when its luminance exceeds this internalized upper-limit luminance.

## 1. Introduction

Most objects in the real world are visible because they reflect light. Some objects however emit light themselves; such self-luminous objects typically have a distinct appearance (e.g., traffic lights visually stand out in a scene). However, any light reaching our retina is indiscriminately encoded by three classes of cone signals regardless of whether the light is reflected from a surface or directly emitted from a light source. Thus, judging whether a given object is self-luminous presents a mathematically underdetermined problem to the visual system. The goal of this study is to reveal how our visual system overcomes this computational challenge and generates a luminous percept.

Self-luminous objects normally have a glowing appearance distinct from the appearance of illuminated surfaces. This qualitative difference was formally introduced as a mode of color appearance (Katz, 1935). The original description finely discriminates various categories, but this study concerns two modes: surface-color mode and aperture-color mode, which respectively correspond to the qualities of color appearance for an illuminated surface and a self-luminous object. Color appearance has mostly been studied in the surface-color mode; only a limited number of studies have investigated the nature of the aperture-color mode (e.g., Uchikawa, Uchikawa & Boynton, 1989).

One common approach is to measure the transition luminance between surface-color mode and aperture-color mode, which is known as the *luminosity threshold.* Past studies have investigated what factors might govern this threshold. In an early study, Ullman (1976) extensively discussed potential determinants of luminosity thresholds: highest intensity in a scene, absolute intensity of the stimulus, local or global contrast, intensity comparison with the average intensity of the scene, and lightness computation that emphasizes a transient intensity change over space while ignoring a gradual intensity change. It was concluded that although each factor plays a role, none of these factors are sufficient to predict luminosity thresholds. Bonato & Gilchrist (1994) reported quantitative observations that an achromatic surface appears luminous when it has roughly 1.7 times the luminance of a surface that would be perceived as white. For chromatic stimuli, it was repeatedly shown that luminosity thresholds were negatively correlated with stimulus purity in a series of studies (Evans, 1959; Evans & Swenholt, 1967; Evans & Swenholt, 1968; Evans & Swenholt, 1969). Speigle & Brainard (1996) measured luminosity thresholds using real colored objects placed under illuminants of different color temperatures. They supported Evans’s consistent observation about the chromaticity-dependent nature of luminosity thresholds and showed that the color of the illuminant also affects luminosity thresholds. More recently, Uchikawa et al. (2001) pointed out that the brightness of colored surfaces rather than their physical luminance is highly correlated with the luminosity thresholds of colored surfaces. These studies well characterized the properties of a test stimulus and of surrounding contexts that have an impact on luminosity thresholds.

One important open question in the field is whether our visual system bases self-luminous judgements purely on heuristics that extract statistics from the external world. Such a strategy is prevalent in many other visual judgements. For example, the famous anchoring theory determines a reference based on simple statistics in a given scene (i.e., the highest luminance in a scene is defined as white) which has been successful in explaining empirical results involving lightness judgements (Gilchrist & Bonato, 1995; Gilchrist et al. 1999). If our visual system takes a heuristic-based strategy, luminosity thresholds should be susceptible to scene content – for example a combination of surface reflectances that happen to be present in a scene. Alternatively, the visual system might additionally use an internal reference for luminosity judgements that is more robust to the variety of available scene contents. For instance, it was shown that our visual system might use statistical regularities about the possible range of surface colors and illuminant colors (Judd et al., 1964) as a prior to solve an ill-posed problem such as color constancy (Maloney & Wandell, 1986). Also, there are suggestions that color contrast and assimilation arise simply from learning statistical regularities in external environments (Lotto & Purves, 2000; Long & Purves, 2003). The success of these prior-based approaches implies a possibility that humans might take a similar strategy when making self-luminosity judgements.

Thus, one primary focus in this study is to reveal whether luminosity thresholds are determined purely based on rigid heuristics that rely on the statistics of external stimuli or whether the visual system additionally uses internal references to make a luminosity judgement. We specifically built a hypothesis based on the latter view: the visual system internalizes the physical gamut of surface colors under various illuminants and refers to this knowledge when judging whether a given surface is self-luminous. This physical gamut of surface colors is visualized by optimal colors (MacAdam, 1935a; MacAdam 1935b), which will be detailed in the General Method section. In short, optimal colors are the colors with the highest luminance that can be produced by reflected light under a given illuminant, for each possible chromaticity. It is assumed that the visual system estimates the illuminant color and chooses the gamut under the estimated illuminant. In a more general sense, this hypothesis could be treated as a Bayesian framework where the visual system monitors the scene illuminant and selects which prior to use based on the estimated illuminant. This hypothesis was specifically designed based on observations made in a series of color constancy experiments (Uchikawa et al, 2012; Fukuda & Uchikawa 2014; Morimoto et al, 2016; Morimoto et al, 2021). In these studies, we developed a model for illuminant estimation that operated on the assumption that the visual system internalizes the gamut of surface colors under various illuminants (i.e., distribution of optimal colors) and the model accounted for observers’ estimations of illuminants reasonably well in a variety of conditions. One interpretation of luminosity thresholds is that the visual system takes the upper-limit boundary of surface colors as the point beyond which objects are self-luminous. Thus, we speculated that the loci of luminosity thresholds measured under different illuminants might resemble the locus of optimal colors.

We note that this study is also built on previous efforts made by Evans (1958) and Speigle and Brainard (1996) for the following reasons. Evans (1958) in part of his analyses first made a comparison between luminosity thresholds and the optimal color locus, which is the primary purpose of this study. Though it was concluded that luminosity thresholds do not well align with optimal color locus, the research used a simple stimulus configuration where a colored surface was presented with a uniform background. Thus, we believe it is worth testing the accountability of the optimal color model under a wider variety of conditions where richer cues to the illuminant are provided. Speigle and Brainard (1996) was the first study to directly suggest that luminosity thresholds are strongly influenced by illuminant color. They further modeled observers’ luminosity thresholds using the upper-limit luminance of physically plausible surfaces in the real world estimated by a linear model that uses basis reflectance functions obtained via a principal component analysis of Munsell papers. The locus obtained via their method corresponds to a practical upper-limit luminance in the real world as opposed to a theoretical upper-limit luminance defined by optimal colors. Nevertheless, we believe that their suggestion shows conceptual similarity to our hypothesis.

In this study, we conducted three experiments to test our hypothesis. In each experiment, we presented a 2-degree circular colored test field surrounded by many overlapping colored circles. We measured luminosity thresholds as a function of test chromaticities. Experiment 1 was designed to test the degree to which luminosity thresholds were influenced by the color statistics of surrounding stimuli, in this case the geometry of the color distribution. In Experiment 2, we tested the effect of the illuminant as well as the shape of the surrounding color distribution to reveal whether the luminosity threshold loci agree with the optimal color locus under different illuminants (3000K, 6500K and 20000K). In Experiment 3, we measured the loci of luminosity thresholds under atypical illuminants (magenta and green) to investigate whether the loci of luminosity thresholds over chromaticities might differ between chromatically typical and atypical illuminants.

## 2. General Method

### 2.1. Computation of physical upper-limit luminance at a given chromaticity

We can compute the theoretical upper-limit luminance at each chromaticity by calculating the chromaticity and the luminance of its optimal colors. Here we provide a basic idea of optimal color, but a more detailed description is available elsewhere (e.g., Uchikawa et al., 2012, Morimoto, et al., 2021). An optimal color is a hypothetical surface having a steep spectral reflectance function as shown in Figures 1 (a) and (b). There are two types (band-pass and band-stop) and they can have only 0% or 100% reflectances. Changing λ_1_ and λ_2_ generates numerous optimal colors (λ_1_ < λ_2_). To give concrete examples we generated three illuminants of black body radiation: 3000K, 6500K and 20000K. Then 7,644 optimal colors were rendered under these illuminants as shown by small dots in Figures 1 (c) and (d). Panel (c) shows L/(L+M) in MacLeod-Boynton (MB) chromaticity diagram (MacLeod & Boynton, 1979) vs luminance distributions. Panel (d) shows log_10_S/(L+M) vs. luminance distributions. To calculate cone excitations, we used the Stockman & Sharpe cone fundamentals (Stockman & Sharpe, 2000).

**Figure 1:**
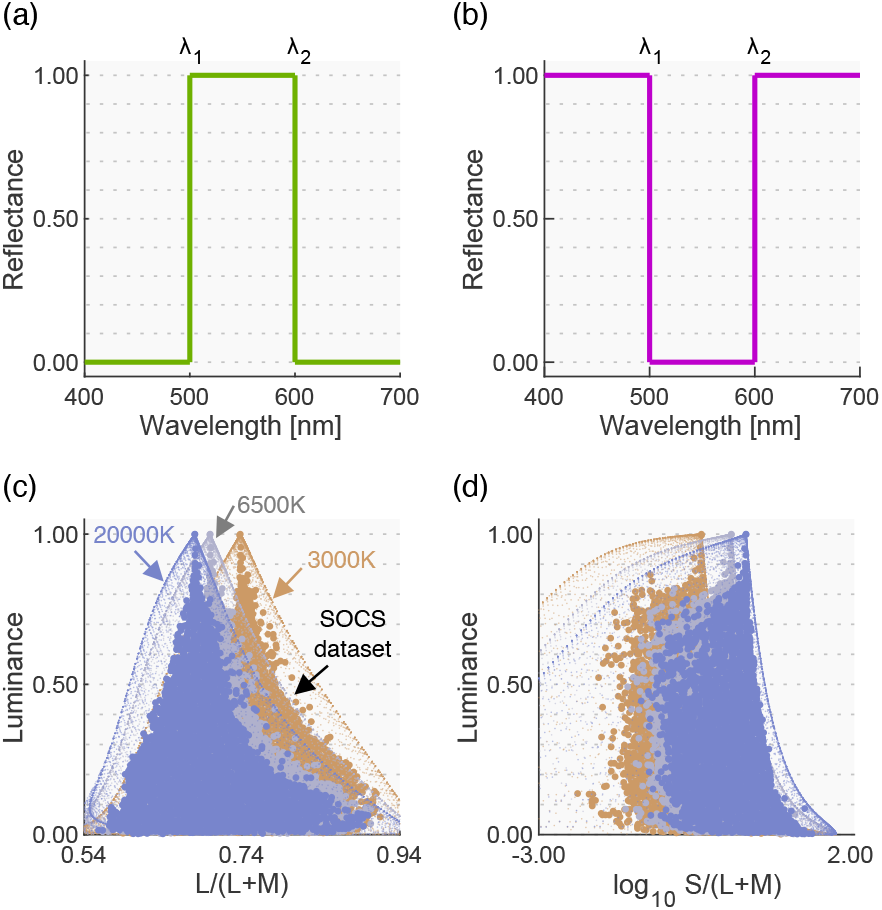
(a), (b) Example optimal colors of band-pass and band-stop types, respectively. (c), (d) L/(L+M) vs. luminance and log10S/(L+M) vs. luminance distributions, respectively, for optimal colors and the SOCS reflectance dataset rendered under 3000K, 6500K and 20000K. [one-column figure]

In the real world surface reflectances must be less than 1.0 at any wavelength due to physical constraints, and thus an optimal color has a higher luminance than any other surface that has the same chromaticity. Thus, no real surface can exceed this optimal-color distribution. To show this concretely, in Figures 1(c) and (d) we plotted 49,667 objects in the standard object color spectra database for color reproduction evaluation (SOCS, ISO/TR 16066:2003).

From optimal color distributions we see that the physical upper-limit luminance is dependent on the chromaticity. The peak of an optimal color distribution always corresponds to a full-white surface (1.0 reflectance across all wavelengths), which thus corresponds to the chromaticity and intensity of the illuminant itself (so-called white point of the illuminant). For this reason, when the color temperature of the illuminant changes, the whole optimal color distribution shifts towards the chromaticity of the illuminant without drastically changing its overall shape. Optimal colors with a higher purity have lower luminance, as they have a narrower-band reflectance, and consequently the distribution spreads out as the purity increases. Importantly, once all optimal colors are calculated, we can look for the physical upper-limit luminance at any chromaticity by looking for the luminance of the optimal color at that chromaticity. Interestingly it is notable that the distribution of real objects (SOCS dataset) shows a somewhat similar shape to the optimal color distribution.

### 2.2. Estimation of the upper-limit luminance at a given chromaticity for real surfaces

The theoretical upper-limit luminance can be computed through the calculation of optimal colors, but the upper-limit luminance for real objects needs to be estimated. Thus, we analyzed 49,672 surface reflectances from the SOCS reflectance database. This dataset includes reflectances from a wide range of categories of natural and man-made objects: “photo” (2304 samples), “graphic” (30,624), “printer” (7856); “paints” (229); “flowers” (148); “leaves” (92); “faces” (8049); and “Krinov datasets” (370) including natural objects which were measured in a separate study (Krinov, 1947). We then excluded reflectances that contained a value higher than 1.0 at any wavelength as they might include fluorescent substances. As a result, one reflectance from the printer category and 4 reflectances from the paints category were excluded.

The remaining 49,667 surfaces were then rendered under 6500K and their chromaticity and luminance were calculated. The luminance value was normalized by that of a full-white surface (100% reflectance at any wavelength). As shown in Figure 2 (a), we plotted the chromaticity of all surfaces on the MacLeod-Boynton chromaticity diagram, where L/(L+M) is the horizontal axis and log_10_ S/(L+M) is the vertical axis. We defined a grid of 25×25 bins and classified 49,667 colors into corresponding bins. Then, for each bin, the maximum luminance across all colors that belong to the bin was defined as the upper-limit luminance of real objects. This procedure was repeated for all 625 bins. The upper-left and lower-left subpanels in panel (b) show the upper-limit luminance for optimal colors (for comparison purposes) and for real objects. As seen here the loci of the upper-limit luminance for real objects were not smooth. We assumed that this is an artifact due to the limited availability of reflectance samples in the database rather than the nature of reflectances of real objects. Thus, we smoothed the upper-limit luminances by spatial filtering with 3×3 convolutional filters (each pixel has the value of 1/9). The lower-right subpanel depicts the smoothed data. Note that this upper-limit luminance heatmap is dependent on the color of the illuminant. Thus we repeated the same procedure for other black-body illuminants with color temperatures from 3000K to 20000K with 500 K steps. Both the optimal color locus and real object locus unsurprisingly peak at the chromaticity of the illuminant shown by the red cross symbol. The upper-limit luminance of real objects decreases more sharply as the stimulus purity increases than that of optimal colors. We can refer to these look-up-tables to find the upper-limit luminance of real objects for an arbitrary chromaticity under illuminants of a range of color temperatures. Note that this upper-limit luminance of real objects corresponds to the proposed model by Speigle and Brainard (1996) at a conceptual level though they estimated the boundary using a linear model rather than the ‘big data’ approach taken here.

**Figure 2:**
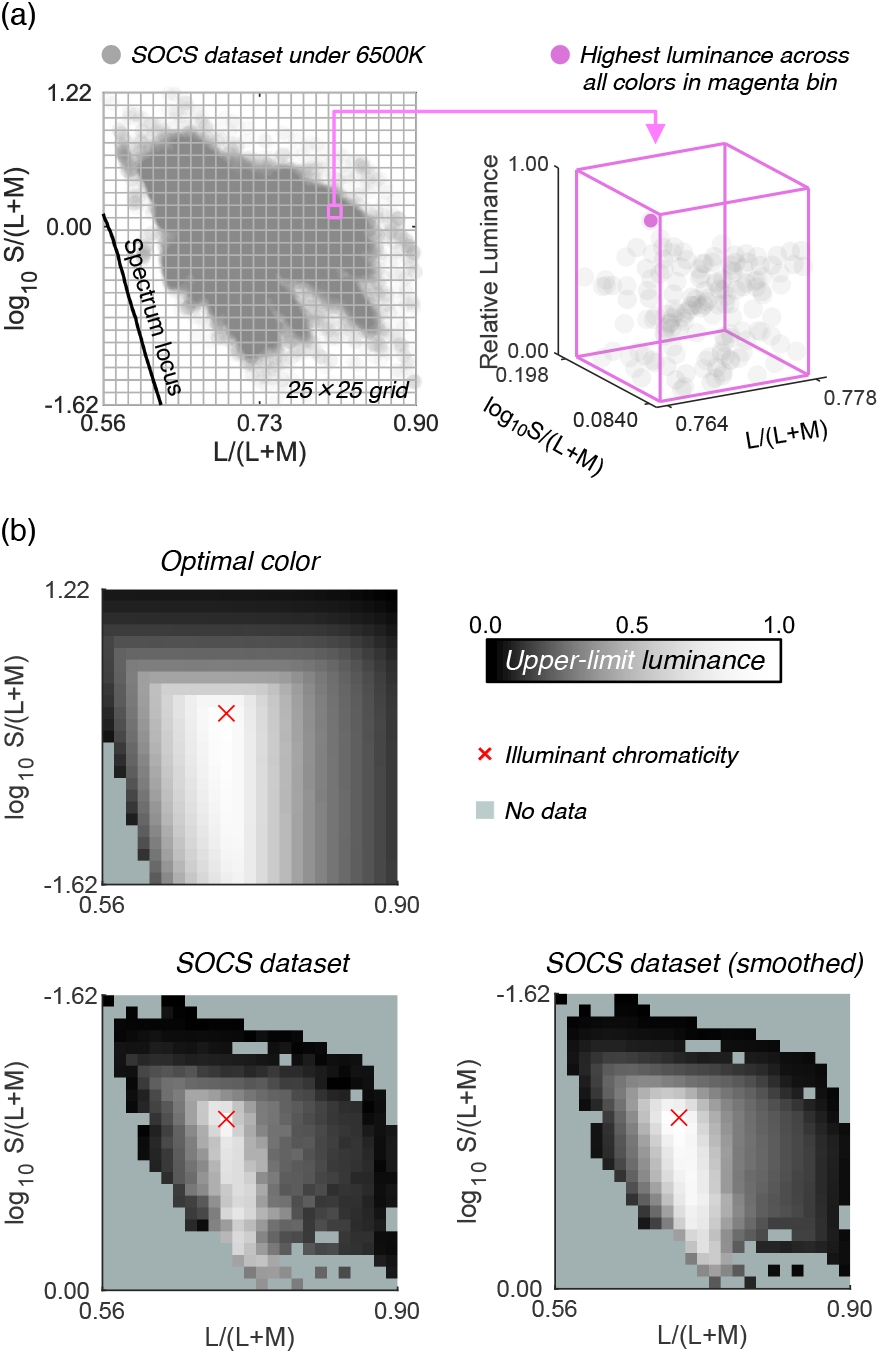
How to estimate the upper-limit luminance for real objects using the SOCS spectral reflectance dataset. A 25×25 grid was first drawn on the MacLeod-Boynton chromaticity diagram. For each grid bin, we searched for the surface that has the highest luminance as shown in the right part of panel (a), which was defined as the upper-limit luminance for that chromaticity bin. Panel (b) shows the locus of upper-limit luminance for optimal color, real objects (raw), and real objects (smoothed) under 6500K illuminant. The lightness indicates the upper-limit luminance value for the chromaticity bins. The pale green color indicates that there is no data in that bin. [one-column figure]

### 2.3. Observers

Four observers (KK, MI, TM and YK) participated in Experiment 1. KK and YK were also recruited for Experiment 2 as well as two new observers (KS and NT). KK, KS and YK participated in Experiment 3. Observers except for KS were naïve to the purpose of all experiments. Observers’ ages ranged between 22 and 57 (*mean* 31.4, *s.d.* 13.2). Observers were all Japanese. All observers had corrected visual acuity and normal color vision as assessed by Ishihara pseudo-isochromatic plates. Before the experiments, informed consent was obtained from each observer. Observers were offered to take several breaks during the experiments, and observers could stop the participation at any point during the experiments.

### 2.4. Stimulus Configuration

The stimulus configuration is shown in Figure 3. The color distribution of the surrounding stimuli and the chromaticities used for the test field are detailed in each experimental section. The spatial pattern was shuffled for each trial.

**Figure 3:**
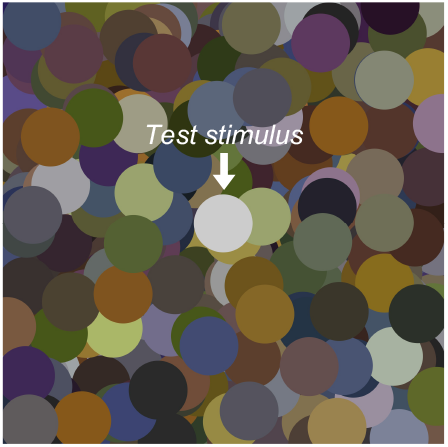
Example of stimulus configuration. The center circle is the test stimulus, and its luminance was adjusted by observers. Each circle had a diameter of 2 degrees in visual angle, and the whole image subtended 15×15 degrees. The surrounding color distribution is detailed in each experimental section. [one-column figure]

### 2.5. Apparatus

Data collection was computer-controlled and all experiments were conducted in a dark room. Stimuli were presented on a cathode ray tube (CRT) monitor (BARCO, Reference Calibrator V. 21 inches, 1844 × 1300 pixels, frame rate 95Hz) controlled with ViSaGe (Cambridge Research Systems), which allows a 14-bit intensity resolution for each of the RGB phosphors. We performed gamma correction using a ColorCAL (Cambridge Research Systems) and spectral calibration was performed with a PR650 spectroradiometer (Photo Research inc.). Observers were positioned 114 cm from the CRT monitor and the viewing distance was maintained with a chin rest. Observers were asked to view the stimuli binocularly.

### 2.6. General procedure

Observers first dark-adapted for 2 mins and then adapted to an adaptation field for 30 seconds. The adaptation field was the full uniform screen that had either a chromaticity of 6500K (Experiments 1 and 3) or the chromaticity of the test illuminant (Experiment 2), and in either case the luminance was equal to the mean luminance value across surrounding stimuli. Then, the first trial began. We drew surrounding stimulus circles so that they had a specific color distribution as detailed in each experimental section. The 2-degree circular test field was presented at the center of the screen. The test field was never occluded by surrounding stimuli. The observers’ task was to adjust the luminance of the test field to the level at which the surface-color mode changed to the aperture-color mode using a keyboard with three possible luminance steps (±0.5, ±1.0, or ±5.0 cd/m^2^). The ambiguity regarding the criterion to judge the transition between surface-color mode and aperture-color mode was reported in a past study (Speigle & Brainard, 1996, Uchikawa et al., 2001). This is mainly because the transition is not sharp, and there is a range that a surface can appear a mixture of surface-color mode and aperture color mode. Considering this reported ambiguity, we instructed observers as follows: “Your task is to adjust the luminance of the center test field so that the test field appears to be at the midpoint between the upper-limit of the surface color mode and the lower-limit of the aperture color mode.” The upper-limit of the surface color mode and the lower-limit of the aperture color mode were described to observers as the limit at which the test field completely appears as an illuminated surface and the limit at which the test field completely appears as a light source, respectively. All observers agreed that this was a reasonable judgment. Also, we note that our criterion is analogous to criteria used in past studies (Bonato & Gilchrist,1994; Evans, 1959; Evans & Swenholt, 1967, 1968, 1969; Speigle & Brainard, 1996; Ullman, 1976). During the experiments, observers were instructed to view the whole stimulus rather than fixate at a specific point to avoid local retinal adaptation. The initial luminance value for the test field was randomly chosen from 2.0, 5.0, 8.0, 11.0, 14.0, 17.0, 20.0, 23.0, 26.0 and 29.0 cd/m^2^. Specific experimental conditions are detailed in each experimental section.

## 3. Experiment 1

### 3.1 Surrounding color distribution, test illuminant and test chromaticity

In a natural scene, the colors of objects tend to cluster around the white point of the illuminant and the density of colors decreases as purity increases. Consequently, the color distribution tends to form a mountain-like shape as shown in Figure 1 (d). The aim of Experiment 1 was to investigate how the loci of luminosity thresholds change when thresholds are measured in a scene that has an atypical color distribution shape. In an extreme case where observers rely purely on internal criteria to judge the self-luminosity of a surface, luminosity thresholds should not change at all regardless of the surrounding color distribution. However, in contrast if observers make a self-luminous judgement using surrounding colors, for example by estimating the upper luminance boundary from the surrounding distribution, luminosity thresholds should largely change depending on the shape of the surrounding color distribution.

Figure 4 (a) shows the five surrounding color distributions used in Experiment 1. The 6500K illuminant on the black-body locus was chosen as the test illuminant in this experiment. We first defined the *natural* color distribution in the upper-left subpanel and then transformed the distribution to generate 4 atypical color distributions (*reverse*, *flat*, *slope+* and *slope-*) in the following ways. First, to construct the *natural* color distribution, we started with a dataset of 574 spectral reflectances of natural objects (Brown, 2003). Out of the 574 reflectances, 516 reflectances were inside the chromaticity gamut of the experimental CRT monitor when rendered under the 6500K test illuminant. All stimuli were presented via a ViSaGe, which had the technical constraint that only 253 colors could be simultaneously presented. Thus, we selected 253 reflectance samples out of 516 reflectances. The reflectance spectra in the Brown dataset were clustered around a white point in a chromaticity diagram; therefore, if we randomly sample from those spectra, it generates a biased distribution with more data points around the white point. Thus the 253 reflectances were selected such that, when rendered under 6500K, they were approximately spatially and uniformly distributed across a chromaticity diagram: L/(L+M) and S/(L+M).

**Figure 4:**
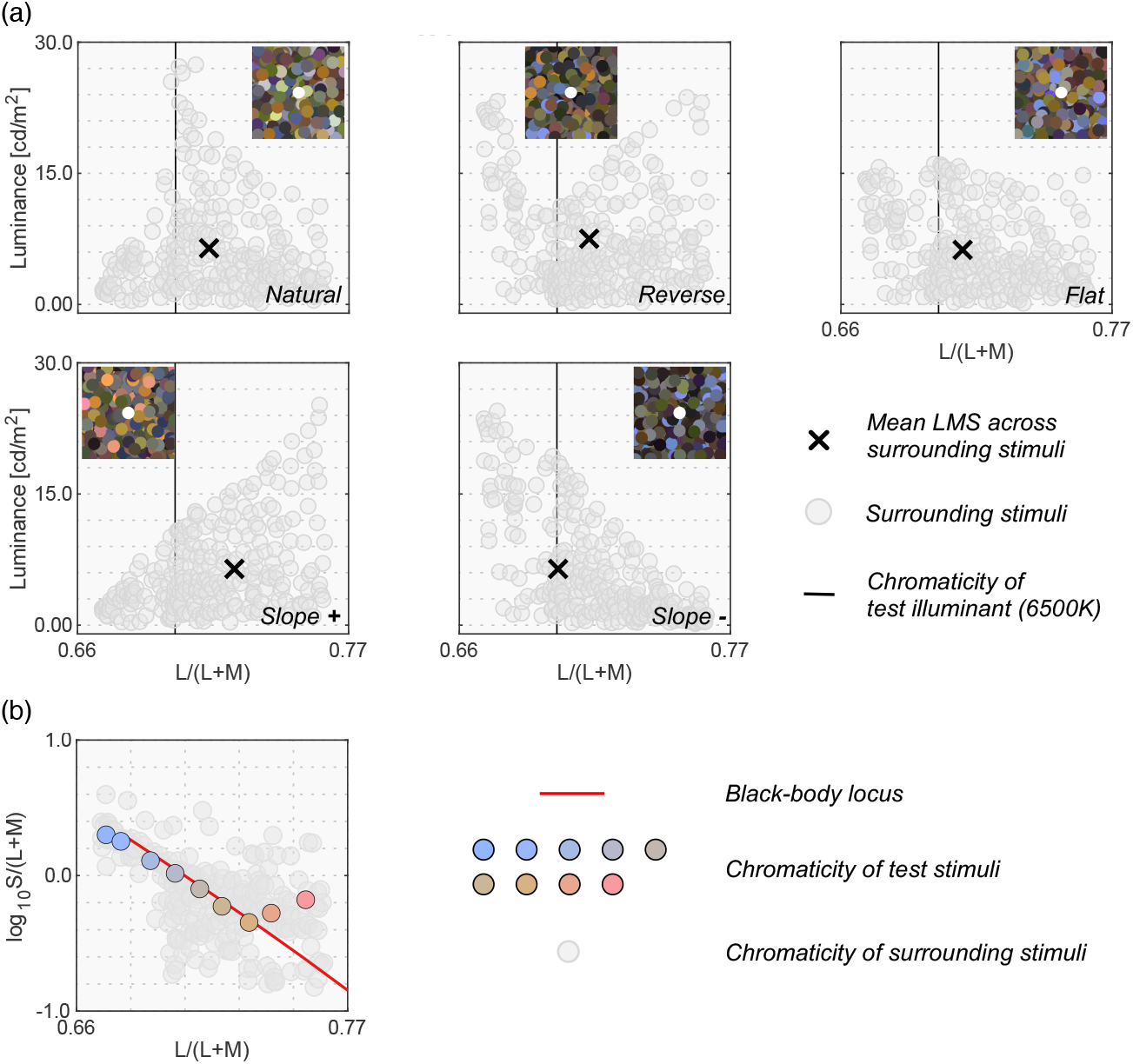
(a) 5 color distribution sets for surrounding stimuli. An inserted image at the top shows an example stimulus configuration for each color distribution. The vertical solid black line indicates the L/(L+M) value of 6500K test illuminant. Black cross symbols indicate mean cone response across all 253 surrounding stimuli. (b) 9 test chromaticities at which the luminosity thresholds were measured. [1.5-column figure]

To generate the other color distributions (*reverse*, *flat*, *slope+* and *slope-*), we independently scaled each of the 253 reflectances by a scalar value to manipulate the luminance while keeping the chromaticity constant. The inserted image in each subpanel shows an example of surrounding stimuli that has the corresponding color distribution. Note that the spatial layout of the surrounding stimuli was shuffled for each trial. For all distributions, the intensity of the test illuminant was determined so that a full-white surface (i.e., 100% reflectance across all visible wavelengths) had a luminance of 35.0 cd/m^2^ under the test illuminant. We note that surrounding colors had relatively low luminance values. For the test field to appear self-luminous, the test field needs to have a substantially higher luminance than the surrounding colors. Thus, the choice of surround luminances was unavoidable in order to ensure that observers could make a satisfactory adjustment at any tested chromaticities within the luminance range allowed by our experimental monitor.

For the center test field, we chose 9 reflectances out of the 253 so that they fell closely along the black-body locus when placed under a 6500K illuminant. The panel (b) in Figure 4 shows these 9 test chromaticities at which luminosity thresholds were measured. The chromaticity of two reflectances are slightly off from the black-body locus. This is because we could not find reflectance samples that exactly fall on the locus.

### 3.3 Procedure

One block consisted of 9 settings to measure thresholds at all 9 test chromaticities in random order. There were 5 blocks in each session to test all 5 distribution shapes. The order of distribution condition was randomized. All observers completed 20 sessions in total (i.e., 20 repetitions for each data point). They completed 10 sessions per day and thus the experiment was conducted in two days.

### 3.3 Results

Figure 5 shows the results for Experiment 1. Colored symbols with error bars indicate each observer’s setting. Each data point is the average across 20 repetitions. The average across 4 observers is shown as black circles. There was some variation across individuals. Furthermore, the experimental design was to try to collect reliable data from a small number of participants. Thus, we discuss results individually. The magenta circles and the line show luminances of optimal colors at test chromaticities when rendered under the test 6500K illuminant (the optimal color locus). In other words, if the visual system uses the optimal color to judge whether a surface emits a light, the observer’s settings should match the magenta line. The blue circles and the line show a smoothed upper-limit luminance locus of real objects, estimated from the SOCS reflectance dataset as shown in Figure 2, which more rapidly decreases as it gets away from the white point than the optimal color locus does. For simplicity, we hereafter refer to the magenta and blue lines as predictions of the optimal color model and the real object model, respectively.

**Figure 5:**
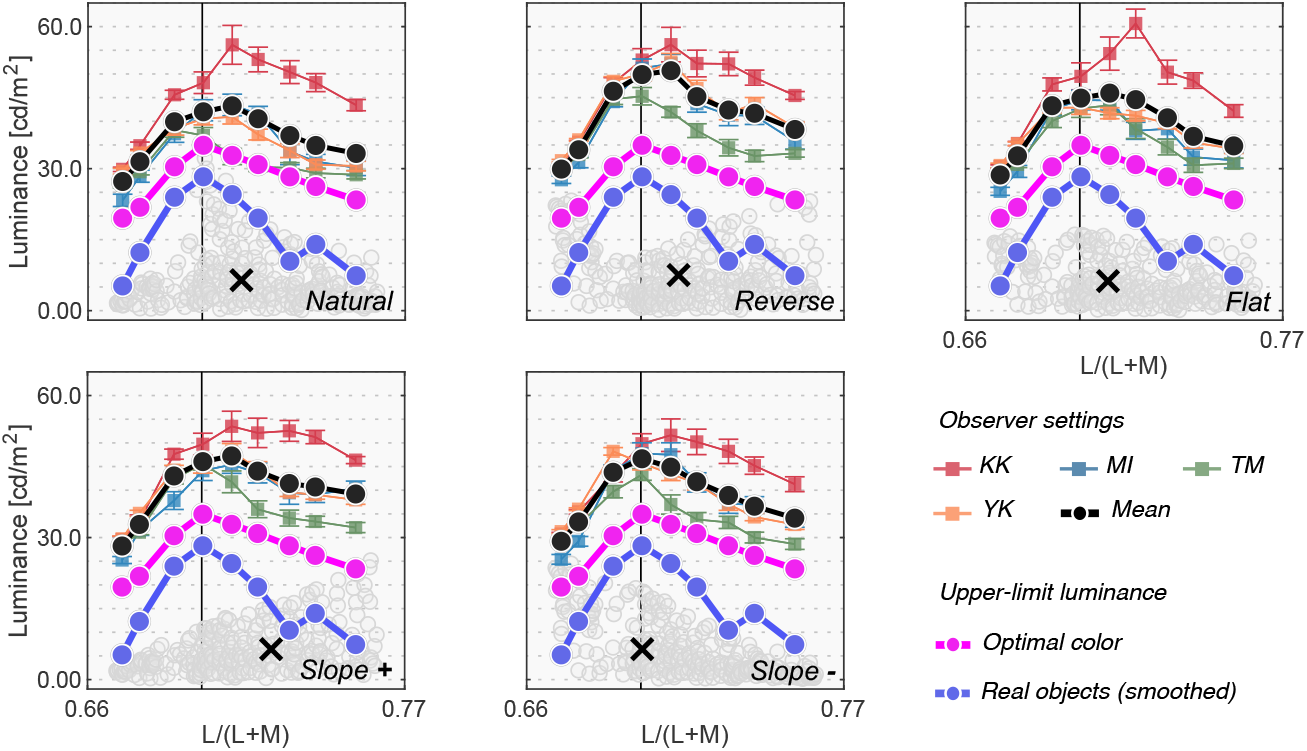
Colored square symbols indicate averaged settings across 20 repetitions for each observer. The error bars represent ±1 S.E. across 20 repetitions. The black circles represent settings averaged across observers (n = 4). The magenta circles denote the luminance of the optimal color at the test chromaticities and thus indicate the physical upper-limit luminance. The blue line shows the upper-limit luminance of real objects estimated from the SOCS reflectance dataset. The vertical solid line shows the chromaticity of the test illuminant (6500K). The black cross symbol indicates the mean LMS value across surrounding stimuli. [1.5-column figure]

First, the loci of luminosity thresholds for all observers had a mountain-like shape regardless of surrounding color distribution. The loci generally peaked around the chromaticity of the test illuminant (the vertical black solid line) and the luminosity thresholds decreased as the test chromaticity moved away from the white point. Although there were some individual differences, especially in the overall setting level (e.g., KK generally had higher thresholds than others) and in the peak chromaticity, the luminosity thresholds generally seem to more resemble the prediction of the optimal color model than that of the real object model in this experiment. This is consistent with the hypothesis that the visual system knows the upper boundary of the optimal color distribution and judges that a given surface is self-luminous when its luminance exceeds the luminance of optimal colors.

To quantify the similarity between observers and models, we calculated Pearson’s correlation coefficient between observer settings and model predictions over the 9 test chromaticities. Figure 6 shows summary matrices of the correlation coefficients. We calculated correlation coefficients for each observer and discuss them on an individual basis.

**Figure 6:**
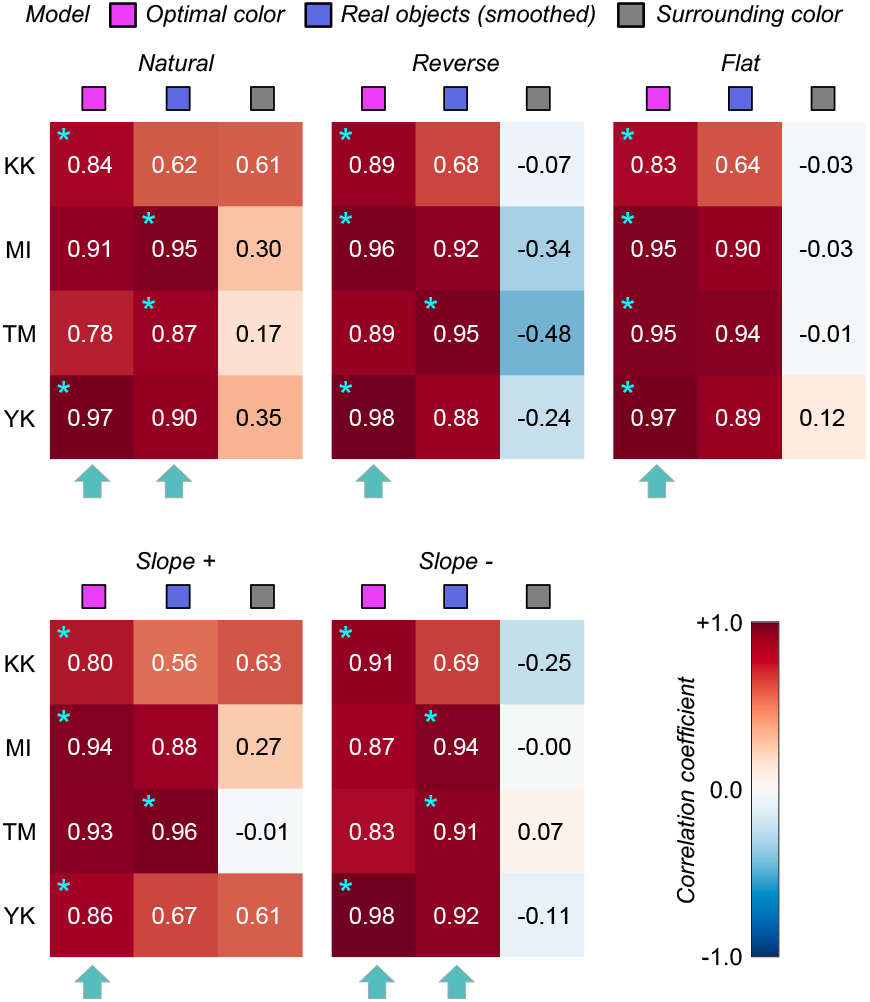
The matrices of Pearson’s correlation coefficients calculated between observer settings and model predictions over 9 test chromaticities. Each subpanel represents each distribution condition (*natural*, *reverse*, *flat*, *slope+* and *slope-*). The color of each individual cell indicates the correlation coefficient as denoted by the color bar. The cyan star symbol indicates the highest correlation coefficient across the 3 models. The cyan arrows at the bottom of each subpanel show the model that received the highest number of cyan star marks, indicating a good candidate model of human observers’ strategy to judge the self-luminosity of a surface. [one-column figure]

The magenta and blue symbols represent the optimal color model and the real object model, respectively. In addition, we evaluated a model which judges the surface as self-luminous when its luminance exceeds that of the surrounding color distribution. The luminosity thresholds estimated from such a model should show much similarity to the shape of the surrounding color distribution. For example, in the *reverse* condition, the luminosity threshold should be lowest at the white point and increase as the saturation of the test stimulus increases. This model is labelled as the “surrounding color” model in Figure 6. Note that this is a simplistic model and we are not trying to claim that the visual system takes such a strategy. Instead, our goal here is to build a framework in which we quantitatively predict an observer’s behavior if she/he judges the luminosity thresholds solely based on surrounding stimuli presented in each trial without using any prior about the statistics of the real world.

The cyan star symbols in some cells indicate the highest correlation-coefficient value across the 3 tested models. The cyan arrows below each subpanel indicate the model that received the highest number of cyan stars across the 4 observers.

Overall, since the observer settings are stable across all distribution conditions, the correlation coefficient patterns are also similar between the optimal color and real object models whose predictions are both not affected by surrounding colors. However, the correlation coefficients for the surrounding color model strongly depend on distribution condition as predicted. Specific trends are as follows. For observers KK and YK, the loci of the luminosity thresholds showed the highest correlation with the optimal color model for all distributions. For TM, the real object model was the best predictor in all distributions except for the *flat* condition. For MI, the optimal color model showed the highest correlation for *reverse*, *flat* and *slope+* conditions while the real object model showed the highest correlation for *natural* and *slope-*conditions. If we summarize these trends based on the number of cyan arrows each model received, the optimal color model is the best predictor in Experiment 1.

The major finding in this experiment is that the loci of luminosity thresholds are nearly invariant regardless of the shape of the surrounding color distribution. This result supports the idea that observers use an optimal color distribution as an internal reference to determine the luminosity thresholds. In the Appendix, we also provide two other alternative models that predicts luminosity thresholds based on post-receptoral signals or cone signals of the test field alone, but it was shown that these models did not predict the luminosity threshold well in Experiment 1. In Experiment 2, we tested whether this observation holds under different illuminants which shift the peak of the optimal color distribution as shown in Figure 1. If the visual system indeed uses optimal colors, changes in luminosity thresholds should reflect changes in the optimal color distribution.

## 4. Experiment 2

### 4.1 Surrounding color distribution, test illuminant and test chromaticity

We employed *natural*, *reverse* and *flat* distributions of surrounding colors. For test illuminants, we used 3000K, 6500K and 20000K on the black-body locus. Out of the 253 reflectances we used in Experiment 1, only 180 samples were inside the chromaticity gamut of the experimental CRT monitor under all test illuminants and those 180 samples were used as surrounding stimuli in Experiment 2. Panel (a) in Figure 7 shows all 9 test surrounding conditions (3 distributions × 3 test illuminants). Though we found that surrounding color distribution has no systematic effects on luminosity thresholds in Experiment 1, we again manipulated the distribution shapes in Experiment 2 to investigate if this finding held under different illuminants.

**Figure 7:**
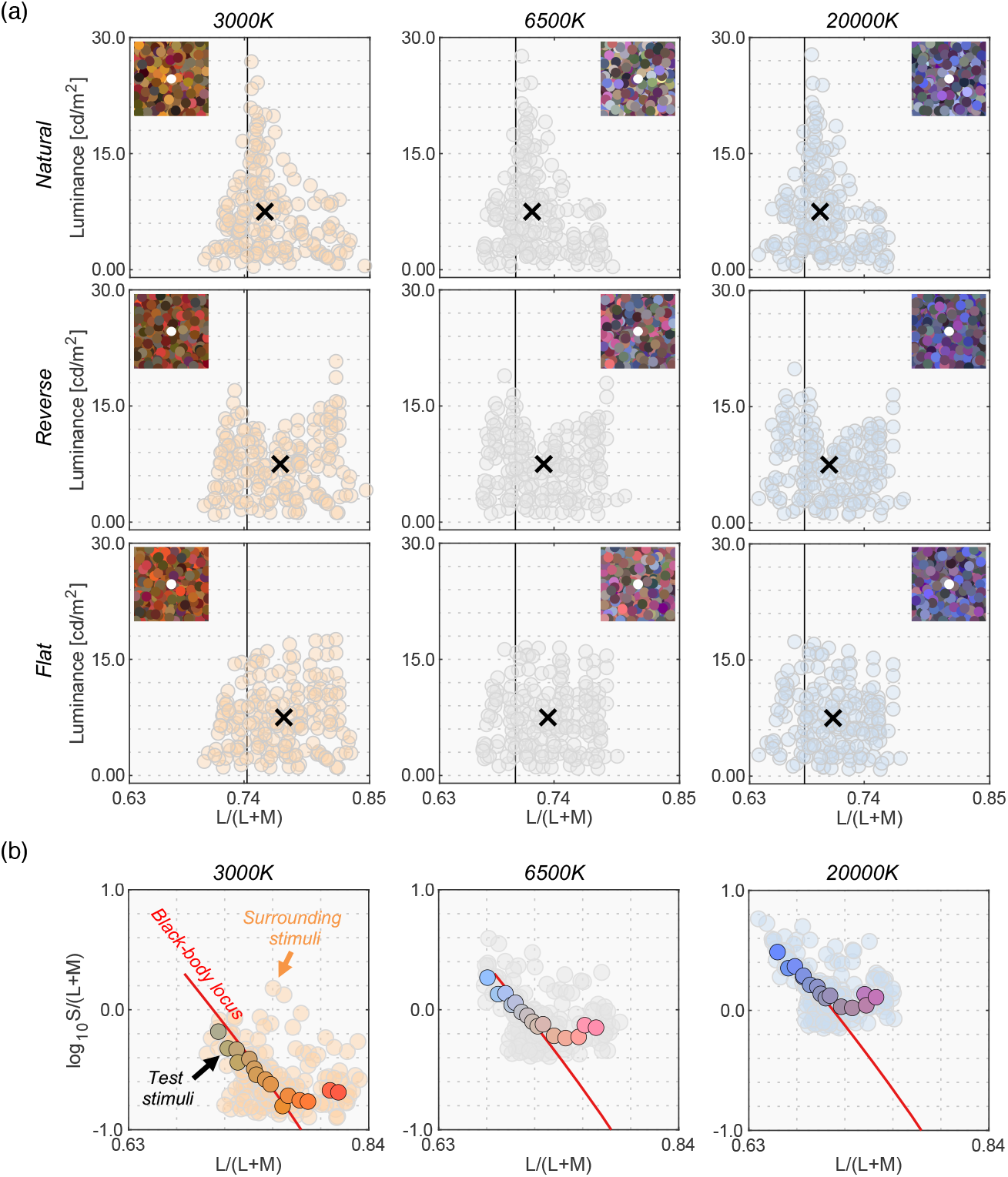
(a) 9 color distributions for surrounding stimuli (3 test illuminants × 3 distributions). Inserted image shows an example stimulus configuration. The vertical solid black line indicates the L/(L+M) value of each test illuminant. Black cross symbols indicate mean cone response values across the 180 surrounding stimuli. (b) 15 test chromaticities at which the luminosity threshold was measured. [1.5-column figure]

We then selected 15 surface reflectances from the 180 reflectances. Panel (b) shows the 15 test chromaticities when rendered under each test illuminant at which the luminosity threshold was measured.

### 4.2 Procedure

One block consisted of 15 consecutive settings to measure thresholds for all test chromaticities presented in random order. There were 9 blocks in one session to test all conditions (3 illuminants × 3 distributions). The order of conditions was randomized. All observers completed 10 sessions in total. The experiment was conducted in three days.

### 4.3 Results

The black line in Figure 8 shows the mean setting across 4 observers. The rest of the data presentation follows the results in Experiment 1. For clarity, only the averaged setting is shown here, but the individual observers’ data is presented in Figure A1 in the Appendix.

**Figure 8:**
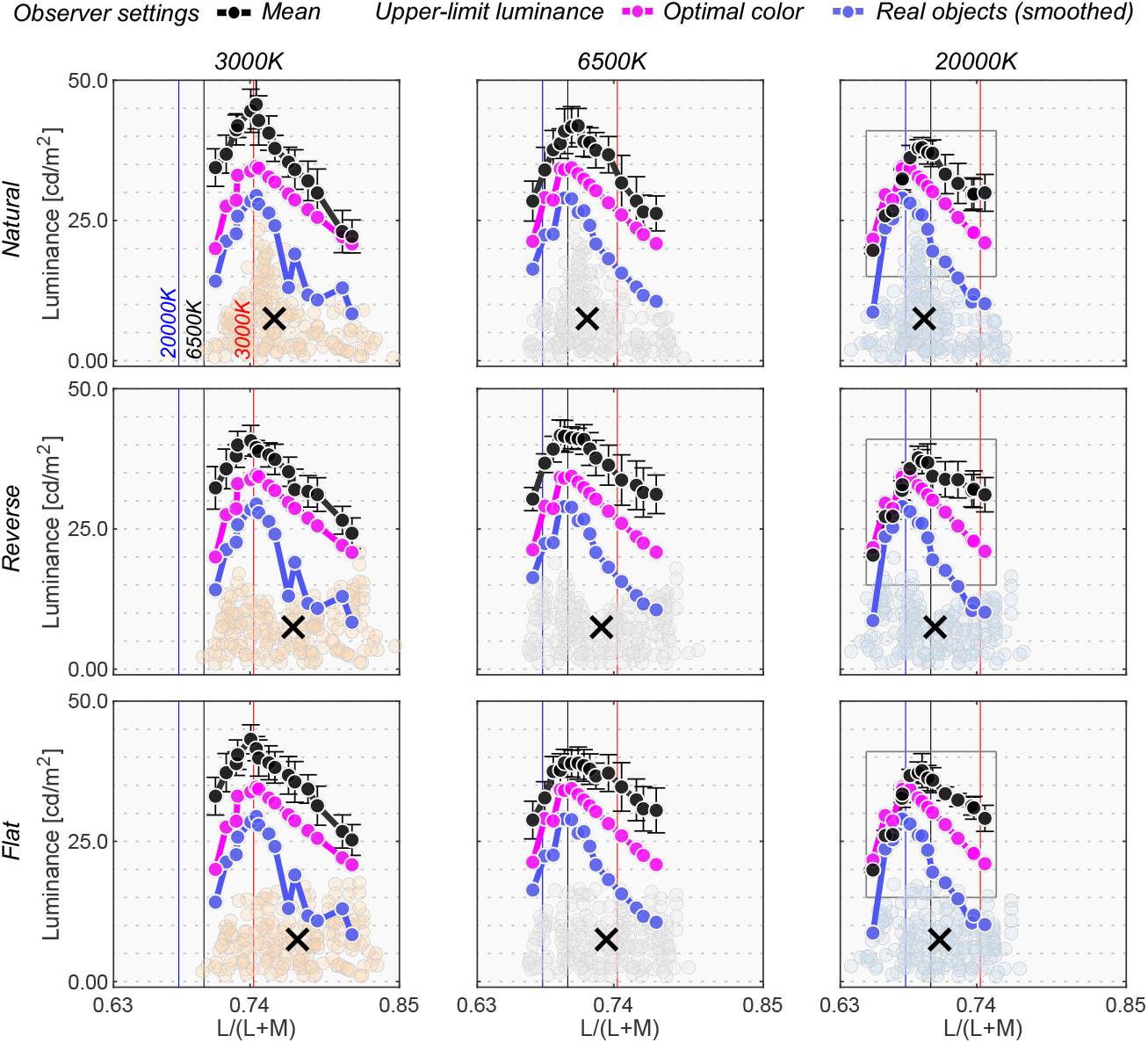
The black circle symbols represent averaged observer settings (n = 4). The error bars represent ±1 S.E. across the 4 observers. The optimal color loci are plotted as magenta circles. The blue circles show the upper-limit luminance of real objects. The red, black and blue vertical solid lines show the chromaticities of the 3000K, 6500K and 20000K test illuminants, respectively. The black cross symbol indicates the mean LMS value across surrounding stimuli. Individual observer data is shown in the Appendix. The region surrounded by a rectangle in the 20000K condition is further discussed in Figure 9. [1.5-column figure]

First, the mean settings showed that the loci of luminosity thresholds were again mountain-like in shape, and the influence of the shape of the surrounding color distribution was almost absent, supporting the findings in Experiment 1. It is also noticeable that the peak chromaticity of the mean setting in each panel shifted towards the illuminant chromaticity shown as vertical solid lines.

It should be noted that the peak chromaticity of the luminosity threshold loci for 20000K was slightly shifted to the right along the L/(L+M) dimension from the chromaticity of the test illuminant. This trend was generally consistent across observers as shown in Figure A1 (Appendix). One potential reason could be that observers misestimated the illuminant color from the surrounding colors. Human color constancy is often imperfect, and thus we speculated that observers’ luminance settings might better agree with the optimal color or real object model rendered under an illuminant estimated by each observer instead of a ground-truth illuminant (20000K). In fact, misestimates of illuminant color was also reported to be an important factor in predicting luminosity thresholds by Speigle and Brainard (1996). The estimated illuminant is typically measured using a technique such as achromatic adjustment (Brainard, 1998), but these data were not collected in this study. Thus, we assumed that the peak chromaticity of observer settings indicated the observer’s estimated illuminant.

We first calculated the chromaticities of illuminants from 3000K to 20000K in 500K steps. Then, for each observer and for each condition independently, we searched for the color temperature that had the closest chromaticity to the peak chromaticity of the luminosity thresholds. Table 1 summarizes the color temperatures of the estimated illuminants in each condition. In the 3000K condition, estimated illuminants matched the ground-truth color temperature for most observers. For 6500K, there was a slight variation across observers. It is notable that in the 20000K condition observers estimated color temperatures substantially lower than those of the ground-truth, meaning illuminant color was estimated to be less blue. This could be because perceptual differences between stimuli rendered under 20000K and under 6500K is smaller than the difference between 3000K and 6500K, as perceptual sensitivities are reported to be worse for bluish illuminants and surfaces (Pearce et al., 2014; Winkler et al., 2015).

**Table 1:** Estimated illuminant by each observer judged from the chromaticity at which luminosity thresholds peaked. The top row shows the color temperatures of ground-truth illuminants, and the other numbers indicate the color temperature of estimated illuminants.

|  |  | 3000K | 6500K | 20000K |
| --- | --- | --- | --- | --- |
| Natural | KK | 3000 | 5500 | 10500 |
|  | KS | 3000 | 7000 | 8500 |
|  | NT | 3000 | 5000 | 10500 |
|  | YK | 3000 | 5500 | 12000 |
| Reverse | KK | 3000 | 6500 | 7500 |
|  | KS | 4000 | 7000 | 7500 |
|  | NT | 3000 | 5500 | 12000 |
|  | YK | 3500 | 5000 | 10500 |
| Flat | KK | 3000 | 7000 | 8500 |
|  | KS | 3000 | 5000 | 8500 |
|  | NT | 3000 | 5500 | 12000 |
|  | YK | 3000 | 6500 | 12000 |

Then, we drew optimal color loci under these estimated color temperatures. This concept is depicted in Figure 9. Intuitively speaking this procedure allows us to estimate an optimal color locus that the observer presumably used during the task, so that the peak of this new optimal color locus coincides with the peak of the measured locus of luminosity thresholds. The cyan curve shows the optimal colors under the estimated illuminant and seems to predict mean observer settings better than the optimal color locus under the ground-truth illuminant (20000K).

**Figure 9:**
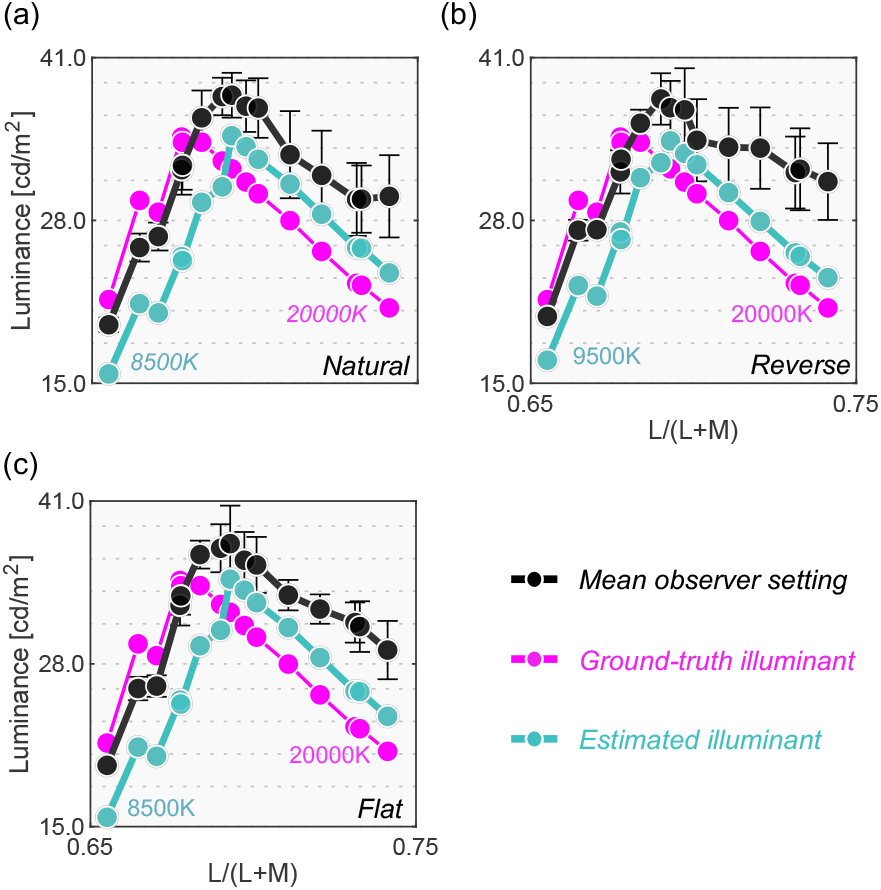
Optimal color models based on the ground-truth illuminant (magenta) and based on estimated illuminants for the averaged observer setting (cyan). It is shown that observers’ settings are better explained by the optimal color model that allows misestimation of illuminants by observers. [one-column figure]

Figure 10 depicts the correlation coefficient matrices for all conditions. We compared correlations from 5 models: (i) the optimal color model and (ii) the real object model under the ground-truth illuminant, (iii) the optimal color model and (iv) the real object model under the estimated illuminant, and (v) the surrounding color model. Again, the cyan star symbol in some cells denotes the highest correlation across the 5 models for that participant. The cyan arrows below each subpanel point to the model that has the highest number of cyan stars – the overall best model for that condition.

**Figure 10:**
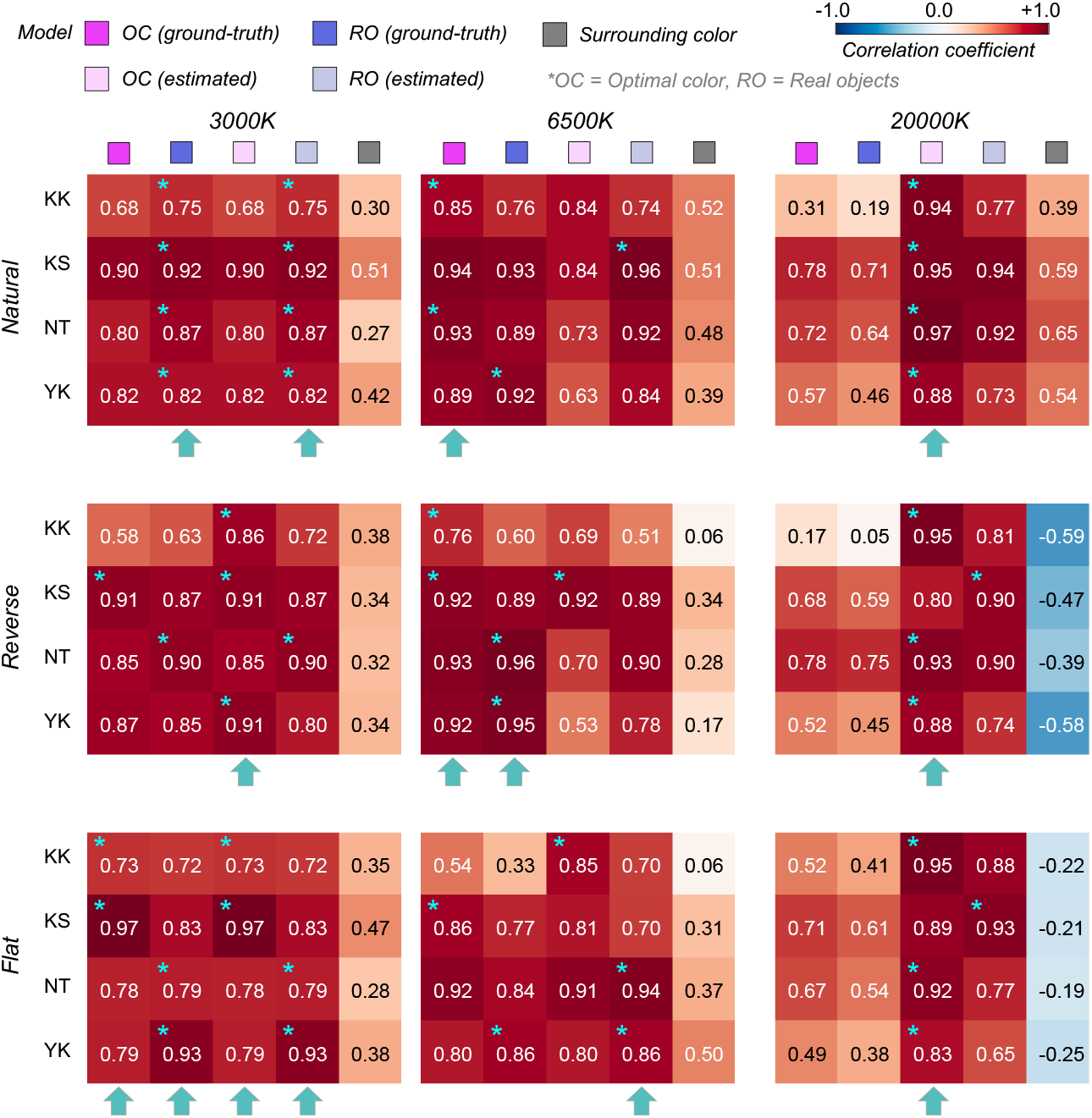
The matrices of Pearson’s correlation coefficient calculated between observer settings and model predictions over 15 test chromaticities. The cyan star symbol indicates the highest correlation coefficient across the 5 models. The cyan arrows at the bottom of each subpanel show the model that received the highest number of cyan star symbols. [1.5-column figure]

Overall, the surrounding color model does not show high correlation with observer settings in any condition, agreeing with the trends in Experiment 1. The optimal color model and the real object model seem to show high correlation, and it depends on the condition which model correlates better. For the *natural-*3000K condition, the highest correlation was found for the real object models, consistently across all observers. For the *reverse*-3000K condition, all observers except NT were best correlated with the optimal color model under the estimated illuminant while for the *flat-*3000K condition the votes were split between the optimal color and real object models. For *natural*-6500K, KK and NT were well predicted by the optimal color model under the ground-truth illuminant, but the other two observers were better correlated with the real object model. For the *reverse*-6500K condition, the optimal color and the real object model both showed high correlations. The real object model, when used with the estimated illuminant, predicted observer settings best for the *flat*-6500K condition. It is notable that for the 20000K condition, the optimal color model under the estimated illuminant was consistently the best predictor. The optimal color model under the ground-truth illuminant also shows much lower correlations, suggesting that observers’ misestimates of the illuminant play a role in predicting luminosity thresholds. In summary both the optimal color model and the real object model showed fairly good agreement with human observers’ settings.

Experiments 1 and 2 collectively suggested that both the optimal color locus and the real object locus seemed to be good candidate determinants of luminosity thresholds. We also note that two other alternative simplistic models again did not predict luminosity thresholds well in Experiment 2 (shown in Appendix). One noteworthy feature in Experiments 1 and 2 is that we used illuminants on the blue-yellow axis that are typically found in natural environments. We also used chromaticities on the black-body locus for the test field. If we assume that the visual system learns the locus of optimal color distribution or real object distribution by observing colors in natural environments, the luminosity thresholds under atypical illuminants may not agree well with the prediction of the optimal color model or the real object model. We directly tested this hypothesis in Experiment 3.

## 5. Experiment 3

Experiment 3 tested whether luminosity thresholds resembled the optimal color locus under atypical illuminants. We also chose a wider range of test chromaticities from the black-body locus and a locus that is orthogonal to the black-body locus.

### 5.1 Surrounding color distribution, test illuminant and test chromaticity

We employed *natural*, *reverse* and *flat* distributions for the surrounding stimuli. For test illuminants, we used magenta and green illuminants. We chose two color filters (Rosco, R44 “Middle Rose” and R4460 “Calcolor 60 Green”) through which the 6500K illuminant was passed to obtain the spectra shown in Figure 11 (a). The chromaticities of these illuminants largely deviate from black-body locus as shown in panel (b). Out of the 574 spectral reflectances of natural objects collected by Brown, 251 reflectances were inside the chromaticity gamut of the CRT monitor under both illuminants. For the surrounding stimuli, we sampled 180 reflectances out of the 251 reflectances and created each distribution following the manipulation used in Experiments 1 and 2. Experiments 1 and 2 showed no effect of surrounding color distribution but Experiment 3 also included this manipulation to confirm that the findings also held under atypical illuminants.

**Figure 11:**
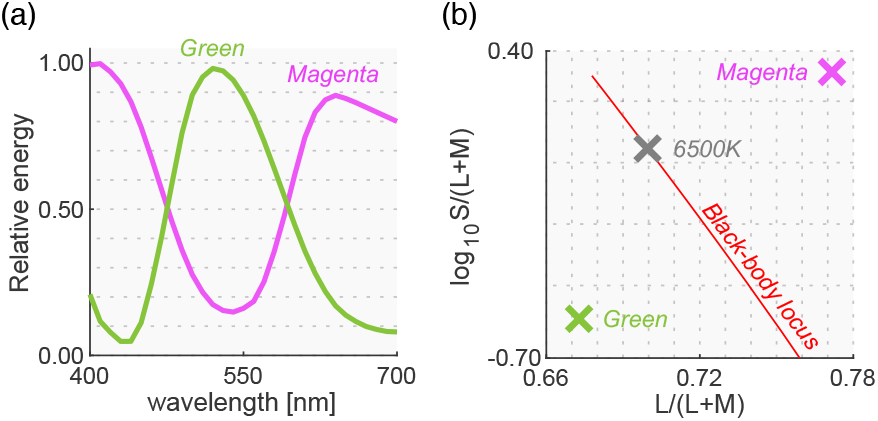
(a) Spectra for magenta and green illuminants used in Experiment 3. (b) Chromaticities of both illuminants. The black-body locus and the chromaticity of 6500K illuminant are shown for comparison purposes. [one-column figure]

Panel (a) in Figure 12 shows surrounding distributions for all 6 test conditions (3 distributions × 2 test illuminants). The intensities of the test illuminants were chosen so that the average luminance across the 180 colors matched 2.5 cd/m^2^.

**Figure 12:**
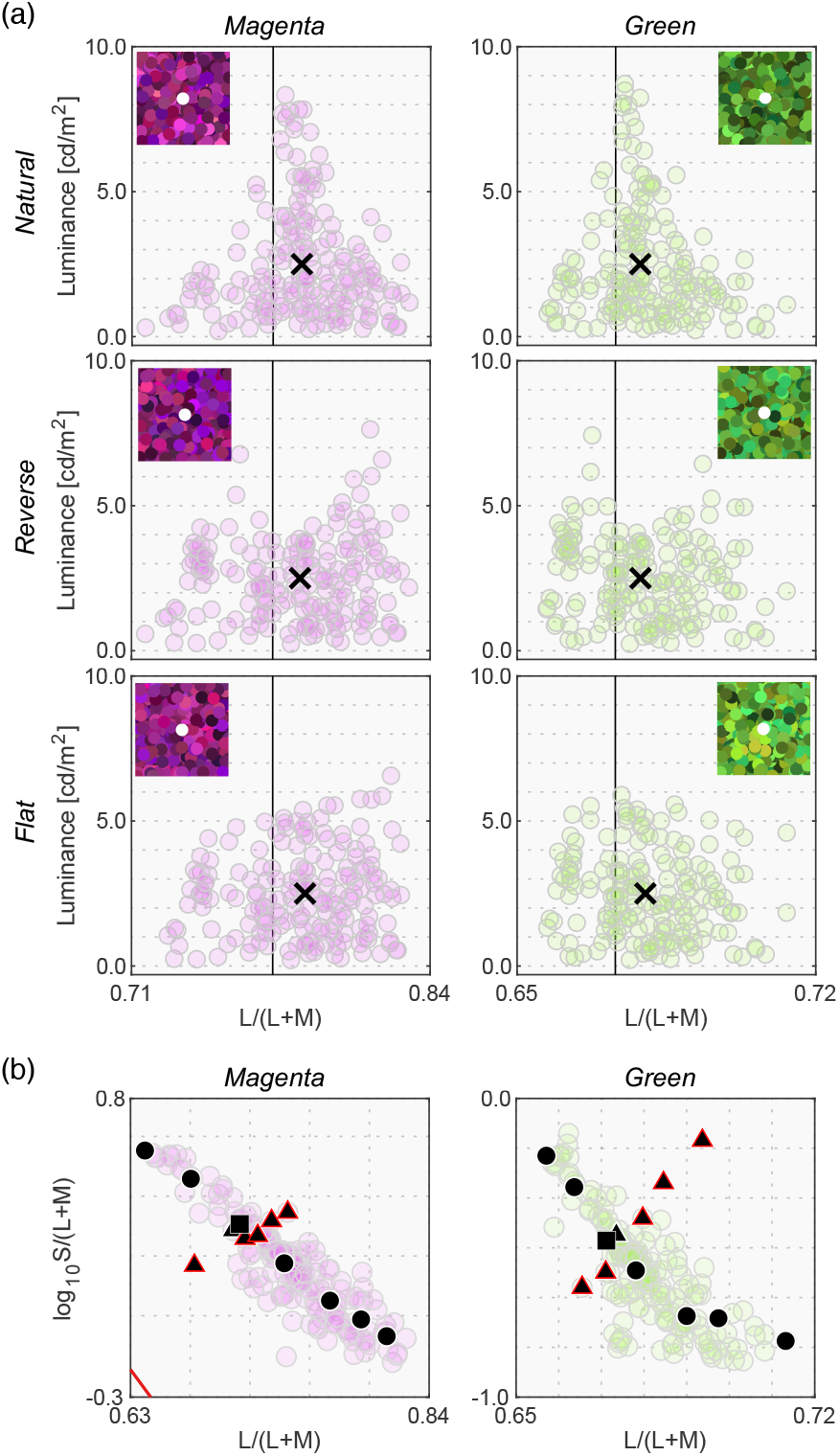
(a) 6 color distributions for surrounding stimuli (2 test illuminants × 3 distributions). Inserted image shows an example stimulus configuration. The vertical solid black line indicates the L/(L+M) value of test illuminant. Black cross symbols indicate mean cone response values across 180 surrounding stimuli. (b) 13 test chromaticities at which the luminosity threshold was measured. Symbols with a red edge indicates reflectances that were not shared between magenta and green illuminants. [one-column figure]

In this experiment, the test chromaticities were chosen so that they varied along two directions: (i) the black-body locus (shown as circles) and (ii) an axis approximately orthogonal to the black-body locus (shown as triangles) depicted in panel (b). First, 8 reflectances were selected from the 180 and were used under both illuminant conditions. Then, we sampled 5 different reflectances separately for each illuminant condition from the reflectances that can be presented only under either magenta or green illuminant. Thus, these reflectance samples are not shared between illuminant conditions. This choice was made to choose test chromaticities on the locus orthogonal to black-body locus as widely as possible. In panel (b), the 5 data points surrounded by a red edge represent the 5 reflectances that were not shared between illuminant conditions. There were 7 chromaticities for each axis, but one chromaticity was used for both axes (plotted as a black square). The chromaticities of natural objects tend to spread along the black-body locus, and the purpose of this design was to test whether luminosity thresholds measured at atypical chromaticities would deviate from the prediction of the optimal color model or the real object model.

### 5.2 Procedure

One block consisted of 13 consecutive settings and thresholds were measured for all test chromaticities in random order. Each session comprised 6 blocks to test all distribution × illuminant conditions. The order of condition was randomized. All observers completed 10 sessions in total. Observers conducted 5 sessions per day and thus the experiment were completed in two days.

### 5.3 Results

Figure 13 shows the results. The left 6 panels depict luminosity thresholds measured at chromaticities along the black-body locus (black circles and square in panel (b), Figure 12) while the right 6 panels indicate thresholds at chromaticities along the orthogonal locus (black triangles and square in panel (b), Figure 12).

**Figure 13:**
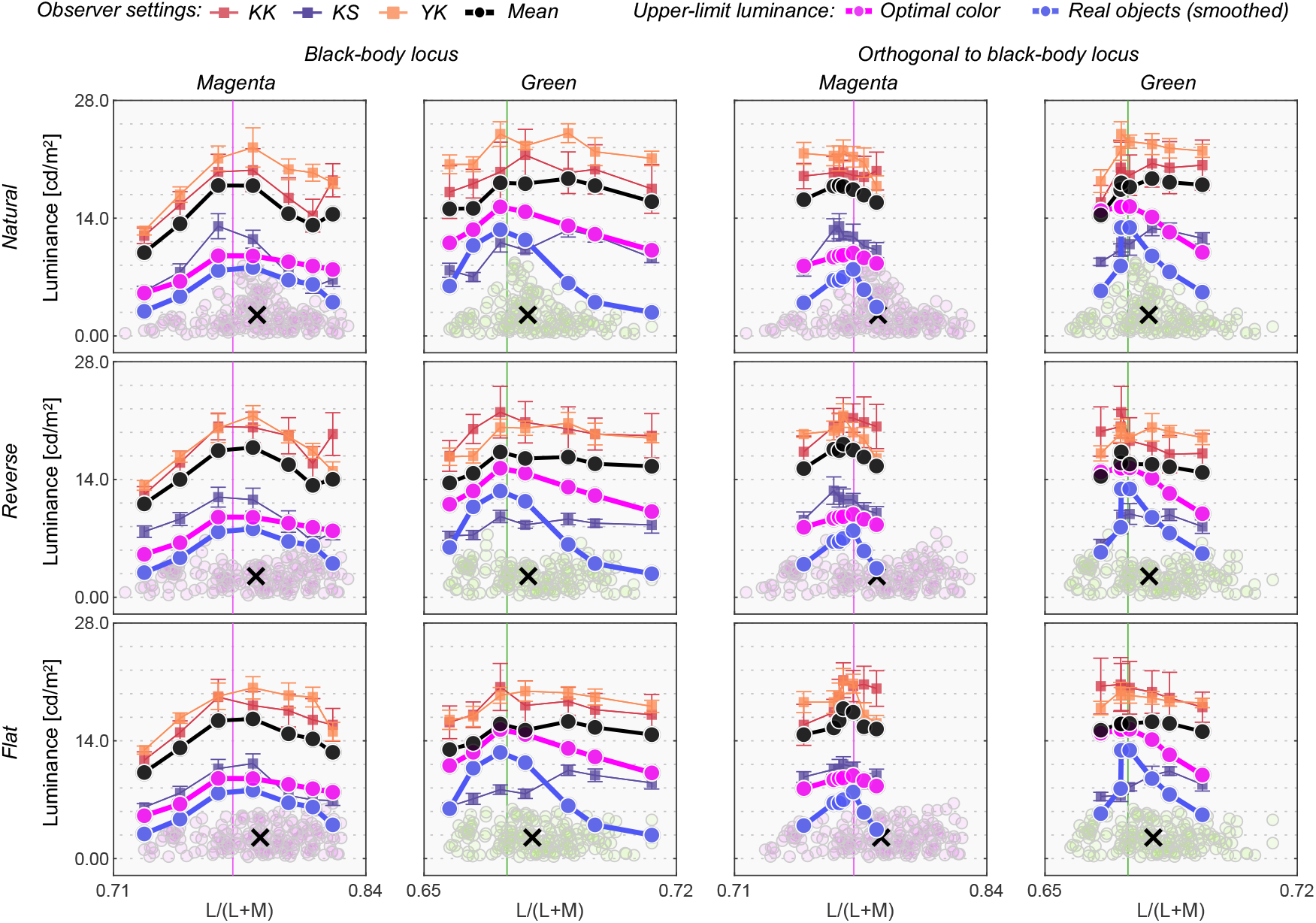
Observer settings in Experiment 3. The left two columns plot luminosity thresholds measured at test chromaticities on the black-body locus (circle and square symbols in panel (b), Figure 12). The right two columns show results for test chromaticities on the locus orthogonal to black-body locus (triangle and square symbols in panel (b), Figure 12). Colored square symbols indicate averaged settings across 10 repetitions for each observer. The error bar plots ±1 S.E across 10 repetitions. The black circle symbols plot average observer settings (n = 3). The magenta circle symbols denote the optimal color locus and the blue circles show the real object locus. The vertical solid line represents the chromaticity of the test illuminant. The black cross symbol indicates the mean LMS value across surrounding stimuli. [two-column figure]

We first look at the left two columns. For the magenta illuminant condition, observers’ settings again show a mountain-like shape. Also, one can see that settings are not dependent on the surrounding color distribution. However, in this condition the optimal color model and the real object model show a relatively flat locus. For the green illuminant, observer settings appear flat. However luminosity thresholds for subject KS show a fairly different trend from the other observers, and the locus is not well predicted by the optimal color locus nor the real object locus, which was not observed in Experiments 1 and 2.

When the test chromaticities are on the axis orthogonal to the black-body locus (right two columns), for the magenta condition all observers’ settings might appear to resemble the optimal color locus. However, for the green illuminant condition, KS again shows a different trend from the other observers and observers do not all agree with either model prediction.

Figure 14 allows us to compare the correlation coefficient across models and conditions. For black-body reflectances shown under the magenta illuminant (the leftmost column), the optimal color model overall showed good correlations for the *natural* condition, while the real object model showed good correlations for the *reverse* and *flat* conditions. For the *natural* condition, one observer (KS, not naïve) had the highest correlation with the surrounding color model, which was not observed in Experiments 1 and 2 in which illuminants on the black-body locus were used as test illuminants. For black-body reflectances shown under the green illuminant (the second leftmost column), in most cells correlation coefficients appeared considerably low. Although the optimal color model consistently had the highest correlation for all distribution conditions (average coefficient across 9 cells is 0.578), the correlation coefficient is not so high if we consider that the correlation for the optimal color model was 0.901 in Experiment 1 (averaged across 5 distributions × 4 observers). Also, in Experiment 2, correlations were 0.746 for the optimal color model of the ground-truth illuminant and 0.837 for the estimated illuminant (average across 9 conditions × 4 observers in both cases).

**Figure 14:**
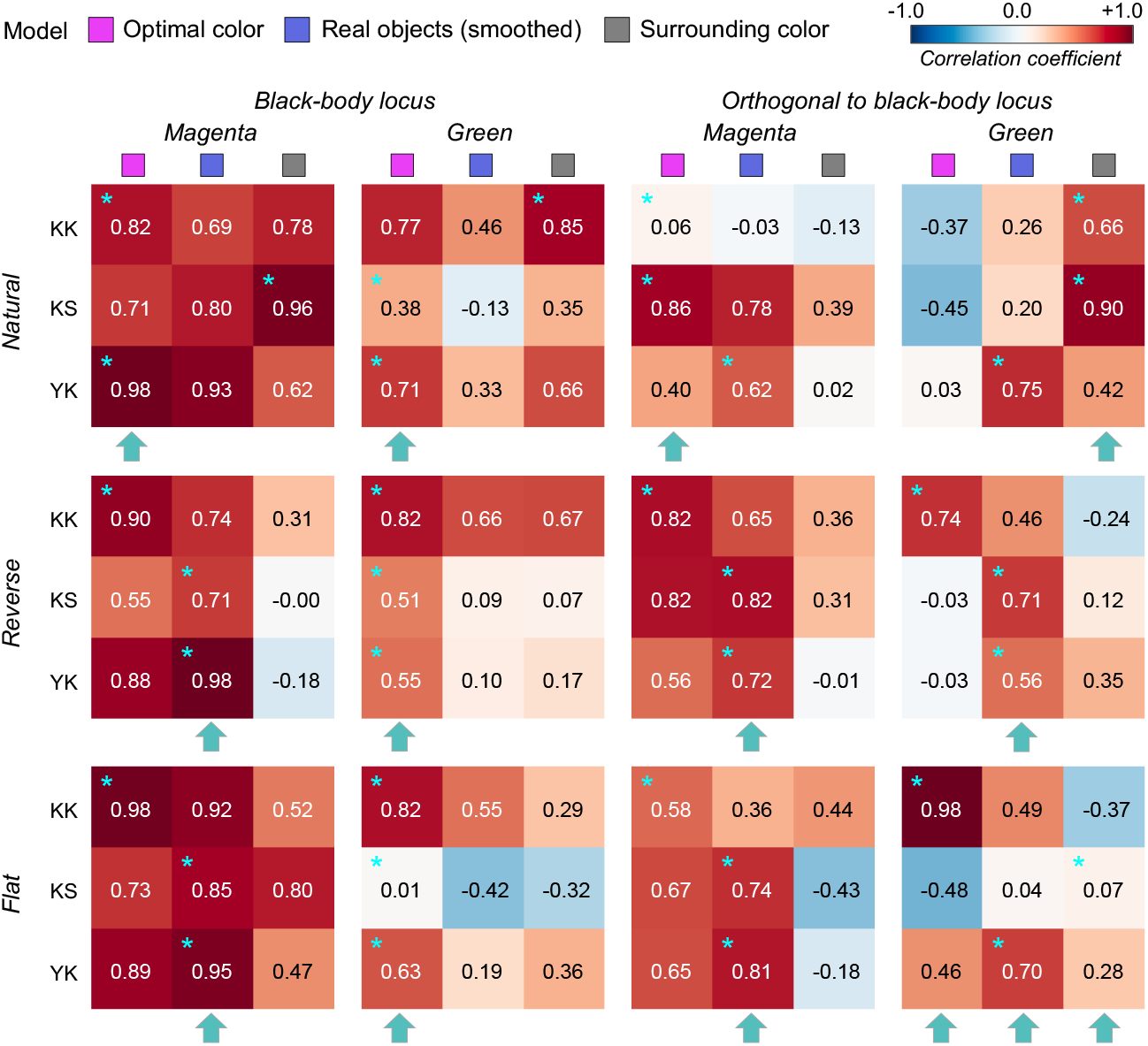
The matrices of Pearson’s correlation coefficient calculated between observer settings and model predictions over 7 test chromaticities in Experiment 3. The cyan star symbols indicate the highest correlation coefficient across the 3 models. The cyan arrows at the bottom of each subpanel indicate the model that received the highest number of cyan stars. [1.5-column figure]

For the reflectances on the axis orthogonal to black-body locus under the magenta illuminant (the second-from-the-right column), the trend seemed to be close to that of the test chromaticities on the black-body locus (leftmost column), but the correlation coefficients overall seemed to be lower. For the green-*natural* condition, the surrounding color model shows a high correlation with observers KK and KS. It is notable that the optimal color model shows nearly zero or even negative correlations. For the *reverse* condition, the real object model showed the best correlation, but their values were not high (0.577, average across 3 observers). For the *flat* condition, we did not find a consistently good model. It may be worth noting that for the green illuminant condition, correlation coefficients for test chromaticities sampled from the locus orthogonal to black-body locus are overall lower than those for test chromaticities sampled from black-body locus.

In summary these results suggested that although the optimal color and real object models can account for observer settings to some extent, overall coefficient values were substantially lower than those observed in Experiments 1 and 2. Also the surrounding color model showed good correlations in some cases. These results might imply that the visual system does not have a rigid internal reference about upper-limit luminance under atypical illuminants and sometimes relies on external cues such as the color of the surrounding stimuli. Also, for green illuminant condition, we found a trend that the predictions of optimal color model were particularly worse when test chromaticities were sampled from the axis orthogonal to the black body locus.

Finally, we summarize results from the three experiments to test whether correlation coefficients of the optimal color model are higher for typical illuminants (Experiments 1 and 2) than atypical illuminants (Experiment 3). For each observer, we averaged the correlation coefficient of the optimal color model across all condition in Experiments 1 and 2 (14 conditions), which served as a summary statistic for typical illuminants. For the 20000K condition in Experiment 2, we used the correlation coefficient value of the optimal color model under the estimated illuminant as it predicted observer settings substantially better than the model under the ground-truth illuminant. We also calculated averaged correlation coefficients across all conditions in Experiment 3 (8 conditions) per observer. The averaged correlation coefficients across all observers were 0.879 ± 0.0114 (average ±1 S.D.) for typical illuminants and 0.525 ± 0.155 for atypical illuminants. Welch’s t-test (one-tailed, no assumption about equal variance) showed that the optimal color model has a significantly higher correlation for typical illuminants than atypical illuminants (t(2.01) = 3.94, p = 0.0290). Also we performed the same analysis using correlation coefficients for the real object model which showed the same trend (t(2.84) = 2.93, p = 0.0326).

These results are consistent with the idea that human observers empirically learn the upper-limit luminance through observing colors in natural environments and use the criterion to judge whether a given surface is self-luminous or not. Since magenta and green illuminants are uncommon in natural environments, the visual system does not know the upper limit of surface colors under those illuminants. Moreover, in the Appendix, we show that a simplistic model predicts the luminosity threshold in Experiment 3 as well as the optimal color model. A potential interpretation would be that when the scene illuminant has an atypical color, the visual system makes a self-luminous judgement based on simple statistics.

## 5. General Discussion

This study investigated potential determinants of luminosity thresholds. Our three experiments showed that the loci of luminosity thresholds have a mountain-like shape that peaks around the illuminant color and decreases as stimulus purity increases, showing a striking similarity to optimal color and real object loci. A simple alternative strategy which bases judgements on the surrounding color distribution did not explain observers’ settings well. Rather observers seem to hold an internal representation about the luminance at which a surface should reach self-luminosity. Moreover, such similarity between luminosity threshold and optimal color or real object loci was higher when surfaces were placed under illuminants along the blue-yellow direction than magenta and green illuminants that are atypical in natural environments. These support an idea that the visual system empirically internalizes the gamut of surface colors through an observation of colors in daily life. Going back to the original question of whether the visual system relies on a heuristic or internal reference for luminosity judgements, the present study generally supports the internal reference hypothesis.

We also note that some properties of the test field we did not consider in the present study are known to affect luminosity judgement. For instance, it was reported that surfaces with smaller areas appeared to emit light at lower luminance levels (Bonato & Gilchrist, 1999). It has also been reported that surround stimuli are more likely to affect luminosity threshold if the surround stimuli are presented at the same depth as the test field (e.g., Yamauchi & Uchikawa, 2005). Thus, in future studies, it would be desired to expand our model to modify the prediction of luminosity thresholds based on factors such as stimulus size and relative depth in a way that agrees with human luminosity judgements.

One consistent trend across the three experiments was that observer settings were above the physical limit (i.e., prediction of the optimal color model) in most cases. We suspect that there are at least two reasons for this. First, observers’ criterion to judge the luminosity threshold in this study was the midpoint between the upper-limit of the surface color mode where a test field purely appears as an illuminated surface and the lower-limit of the aperture color mode where a test field purely appears as a light source. If we instead used a different criterion, for example to set the luminance so that the test field appears simply as the upper-limit of the surface color mode, we would have seen a smaller discrepancy. Second, it is possible that observers overestimated the intensity of the test illuminant. Just as humans cannot directly access the chromaticity of test illuminant, the intensity is also not directly known to observers. In fact, in other color constancy studies (Morimoto, Fukuda & Uchikawa 2016; Morimoto et al., 2021), we have repeatedly found that observers tend to overestimate illuminant intensities and the degree of overestimation substantially varied across individuals. When the illuminant intensity is overestimated, the observers’ internal upper-limit luminance should accordingly increase, which could account for an observed discrepancy between model predictions and observer settings. Moreover, we also note that the limit drawn by optimal-color or real-object model corresponds to the upper-boundary as a surface color, but the models are not designed to exactly predict how the test field should appear when the luminance of the test field exceeds the predicted value by the models. Thus, it is inherently difficult to directly compare the model prediction and observers’ settings, and we think that they rather need to be compared relatively (for example, using correlation coefficient). We also found that individual differences in this study were mainly found in gain rather than the shape of the observer settings (though for some conditions in Experiment 3 shape differences were also evident). This individual variation in gain could also be due to the variability in estimating illuminant intensity.

Color constancy is often described as a visual ability to identify the same surface under different illuminants. A surface reflects a light, and that reflected light enters our eyes. Because the reflected light is a product of surface and illuminant components, color constancy is often framed as a process in which our visual system estimates the influence of the illuminant. The “brightest is white” heuristic, which assumes that a surface with the highest luminance provides the closest information about the illuminant color, has been known as an influential approach in estimating illuminant color (Land, 1977). However, self-luminous objects do not carry information about the scene illuminant, which might cause a misestimation of the illuminant if included in a scene. In general, when we receive an intense light from a surface, there are two ways to interpret this. One is that the surface is placed under an intense illuminant and the other is that the surface is self-luminous. This example highlights the need for luminous percepts to be incorporated into the process of color constancy. In fact, Fukuda & Uchikawa (2014) showed that a surface appearing in aperture-color mode does not have a strong influence on observers’ estimates of the illuminant.

We chose a set of colored circles as experimental stimuli to directly test our hypothesis while excluding any other cues. However, it is reported that changing a material property could affect the mode of color appearance (Kuriki, 2015). Also, our experimental stimuli were simulated to be uniformly illuminated by a single illuminant, but in natural environments the spectra hitting an object surface changes from one direction to another (Morimoto et al., 2019). The presence of multiple illuminants means that we need to consider multiple optimal color distributions, and thus the loci of luminosity thresholds measured under such an environment might also change. Despite a growing amount of research on material perception (Fleming, 2013), luminosity perception is little studied in the field. While our choice of stimuli was necessary for experimental control, it will be interesting whether our finding can be applied to a wider range of stimuli that have complex material properties and are illuminated in non-uniform ways.

One closely related phenomenon to self-luminous perception would be brightness perception of colored objects. The Helmholtz-Kohlrausch effect is that stimuli with high purity appear to have high brightness even if luminance is kept the same. The effect was reported under a variety of viewing conditions (Nayatani et al., 1991; Donoforio, 2011). However, it is unclear why a color with high purity appears brighter. Curiously, as observed in the present study, the same trend holds for luminosity thresholds: a surface with high purity reaches the limit of surface color mode at a lower luminance level. Thus, if we take a strategy to determine the brightness of colored stimuli in comparison to the theoretical upper-limit luminance at the chromaticity we could account for the Helmholtz-Kohlrausch effect. Uchikawa et al. (2001) directly focused on this relationship and argued that saturated colors appear brighter because the visual system knows that it has a lower limit and brightness might be determined in proportion to the theoretical upper-limit luminance.

Identifying the range of natural colors has been a major focus especially in the field of color science (e.g. Pointer, 1980). While the limit of chromaticity has been well characterized, less is known regarding the luminance limit. In this study, we used the SOCS reflectance dataset as a reference to draw an upper-luminance boundary for real objects. The database covers a wide range of color space as it includes man-made materials such as ink which can have narrow-band reflectances. We do not intend to claim that the SOCS dataset in any sense represents all plausible natural reflectance spectra. Yet, our separate analysis based on 16 hyperspectral images (Nascimento, 2002; Foster, 2006) showed that colors in those images were mostly covered in the gamut of the SOCS dataset. Also, to our knowledge we have not encountered another dataset that has a larger color gamut than the SOCS dataset. We also found that if we restrict samples to natural objects, the color gamut largely shrinks (see Figure 2 (b) in Morimoto et al., 2016), and upper-limit luminance estimated only from natural samples would not predict obtained luminosity thresholds in this study. Also, in this study, we used a smoothed upper-limit luminance. We confirmed that if we instead used raw unsmoothed data, the correlation coefficient was lower in almost all tested conditions. These results show that a precise evaluation of the abundance of reflectance samples in real world seems to play a key role in understanding the luminosity percept. When more reflectance datasets become available in the future, the gamut of real objects may need to be re-evaluated.

In summary, our results showed a mysterious similarity between luminosity thresholds and optimal colors. Yet it is difficult to make a conclusive statement as to whether the optimal color model is better in accounting for luminosity thresholds than the real object model. This is partially because the optimal color locus well resembles the locus of real objects, leading to high correlation between predictions from two models. Furthermore, an intrinsically more challenging question would be how our visual system learns the optimal color locus because optimal colors do not exist in the real world. Considering this point, one plausible theory would be that our visual system learns the plausible range of surface colors by seeing colors in daily life and empirically internalizes the gamut of surface colors. Then, a given surface appears self-luminous when its luminance exceeds the upper-limit luminance empirically internalized in the visual system. This study presents a potential link between our perceptual judgment and statistical properties of the real world.

## Acknowledgement

All authors thank Laysa Hedjar for her careful edit of the language throughout the manuscript. This work was supported by JSPS KAKENHI Grant Number JP19K22881, JP17K04503 and 26780413. TM is supported by a Sir Henry Wellcome Postdoctoral Fellowship awarded from the Wellcome Trust (218657/Z/19/Z) and the Junior Research Fellowship from Pembroke College, University of Oxford. This research was funded in whole, or in part, by the Wellcome Trust (218657/Z/19/Z). For the purpose of open access, the author has applied a CC BY public copyright licence to any Author Accepted Manuscript version arising from this submission.

## Data access

The raw experimental data are available in a data repository at https://doi.org/10.5281/zenodo.5590120. Codes to reproduce figures in the main manuscript are available at https://github.com/takuma929.

## Appendix

### Individual observer settings in Experiment 2

In the main text, we presented mean observer settings for Experiment 2. Figure A1 shows the individual observer settings. There is some individual variation, but overall the trend was similar across individuals.

**Figure A1:**
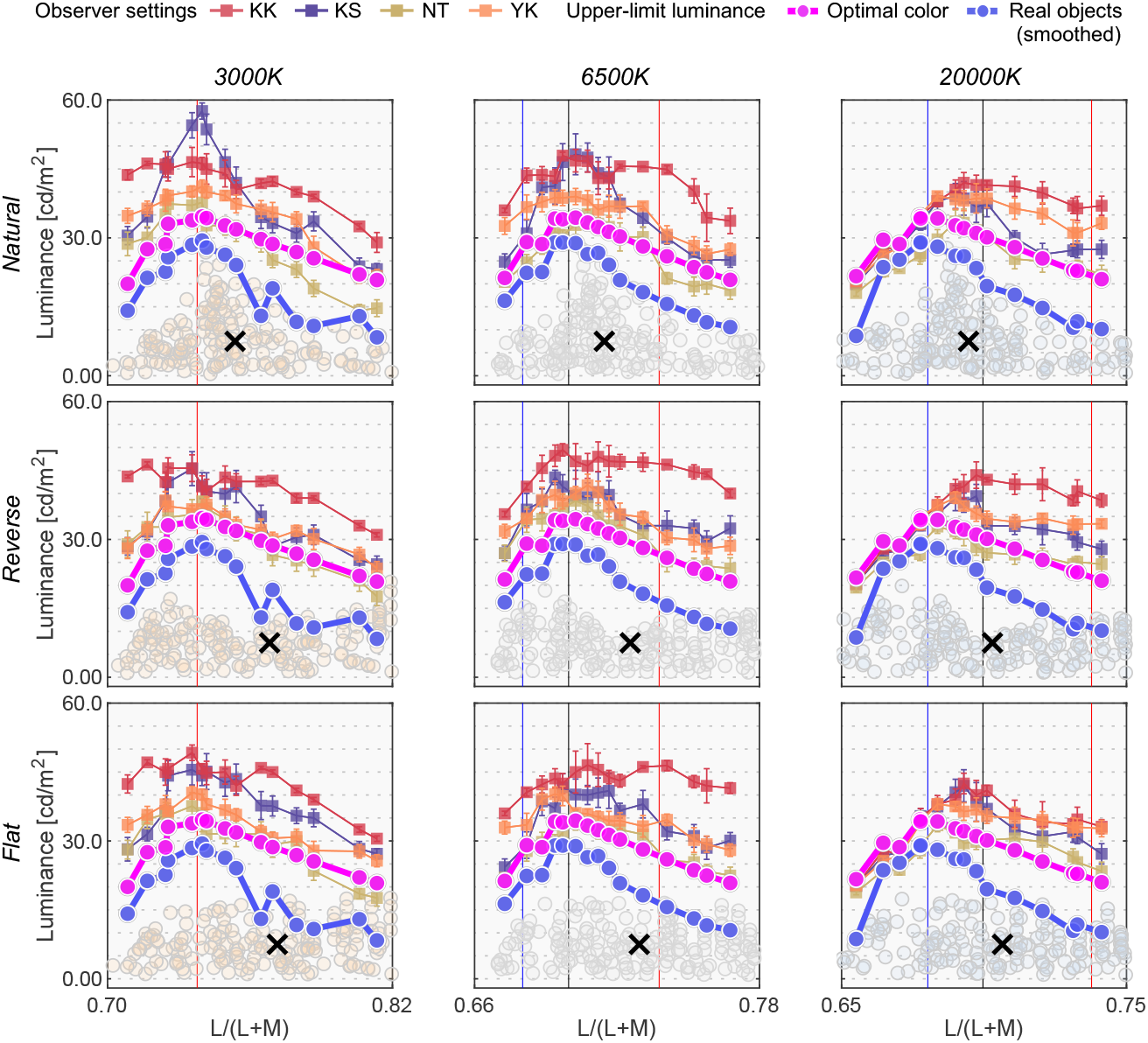
Individual observer settings in Experiment 2. Colored square symbols indicate the averaged setting across 10 repetitions for each observer. The error bar indicates ±1 S.E across 10 repetitions. The magenta circles denote the optimal color locus and the blue circles show the real object locus. The red, black and blue vertical solid lines show the chromaticities of the 3000K, 6500K and 20000K test illuminants, respectively. The black cross symbol indicates mean LMS value across surrounding stimuli. Note that the horizontal range differs across panels. [two-column figure]

### Implementation of other past models

Here, we compare two simplistic models that predict luminosity thresholds based on post-receptoral signals or cone signals of the test field. Note that these two models were also implemented as candidate models in Speigle & Brainard (1996).

The first model is a post-receptoral model which assumes that the visual system monitors the weighted sum of the three types of cone signals of the test field. The test field appears self-luminous when *w_L_L_T_* + *w_M_M_T_* + *w_S_S_T_* >= 1, where L_T_, M_T_ and S_T_ denote cone signals of the test field and *w_L_*, *w_M_* and *w_S_* denote weightings for each class of cone signal (*w_L_* > 0, *w_M_* > 0, and *w_S_* > 0). Then, we optimized weightings *w_L_*, *w_M_* and *w_S_* which produced the minimum root mean square error between model prediction and mean observers’ setting.

The second model is a generalized Evans’s model (Evans, 1959) which assumes that a test field appears self-luminous when any of *L_T_*, *M_T_* or *S_T_* cone signals exceed a certain criterion level. Thus, the model predicts that a test field is self-luminous when *L_T_* >= *c_L_, M_T_* >= *c_M_* or *S_T_* >= *c_S_.* The goal of the optimization here was to find criterion *c_L_*, *c_M_* and *c_S_* that minimized the root mean square error between model prediction and mean observers’ setting.

For both models, the optimization procedure was performed separately for each experimental condition. In other words, for example in Experiment 2, we performed optimization procedures 9 times in total (3 illuminants×3 distributions).

Figures A2, A3 and A4 depict the prediction of the post-receptoral model and the generalized Evans’s model as well as the optimal color model in Experiments 1, 2 and 3. To obtain the prediction of the optimal color model, we used the ground-truth illuminant for Experiments 1 and 3, but for Experiment 2, we used the estimated illuminant so that the peak is matched between the prediction of optimal color model and the mean observer settings (for more details, see the Results section in Experiment 2 in the main text).

For Experiment 1, we see that the post-receptoral model predicts a linear luminosity threshold locus over L/(L+M) which did not lead to a high correlation coefficient (shown at the right upper corner in each panel). The generalized Evans’s model came closer to observer settings, but in all conditions the correlation coefficient was lower than that of the optimal color model. Welch’s t-test (one-tailed, no assumption about equal variance) on averaged correlation coefficients across the 5 conditions showed that the optimal color model has a significantly higher correlation than the Evans’ model (*t*(4.64) = 4.57, *p* = 0.0072).

A quite similar trend is shown in Experiment 2 though in one condition (*flat* and 20000K), the Evans’s model exceeded the optimal color model’s correlation coefficient. However, again Welch’s t-test on averaged correlation coefficients across the 9 conditions showed a significantly higher correlation coefficient for the optimal color model than the Evans’s model (*t*(8.95) = 3.72, *p* = 0.0048).

In contrast, for Experiment 3, we see that the Evans’s model showed higher correlations than the optimal color model especially in green illuminant conditions. Welch’s t-test on averaged correlation coefficients across the 12 conditions showed that there is no significant difference between the Evans’s model and the optimal color model (*t*(12.2) = 1.80, *p* = 0.0962).

In summary, it is evident that for Experiments 1 and 2 the optimal color model predicts human observer settings better than the two alternative models considered here. In Experiment 3, there was no significant difference in correlation coefficients between the Evans’s model and the optimal color model. One interpretation would be that when the scene illuminant is atypical, human observers rely on the simple statistics such as cone signals because the visual system does not know the optimal color locus under the atypical illuminant.

**Figure A2:**
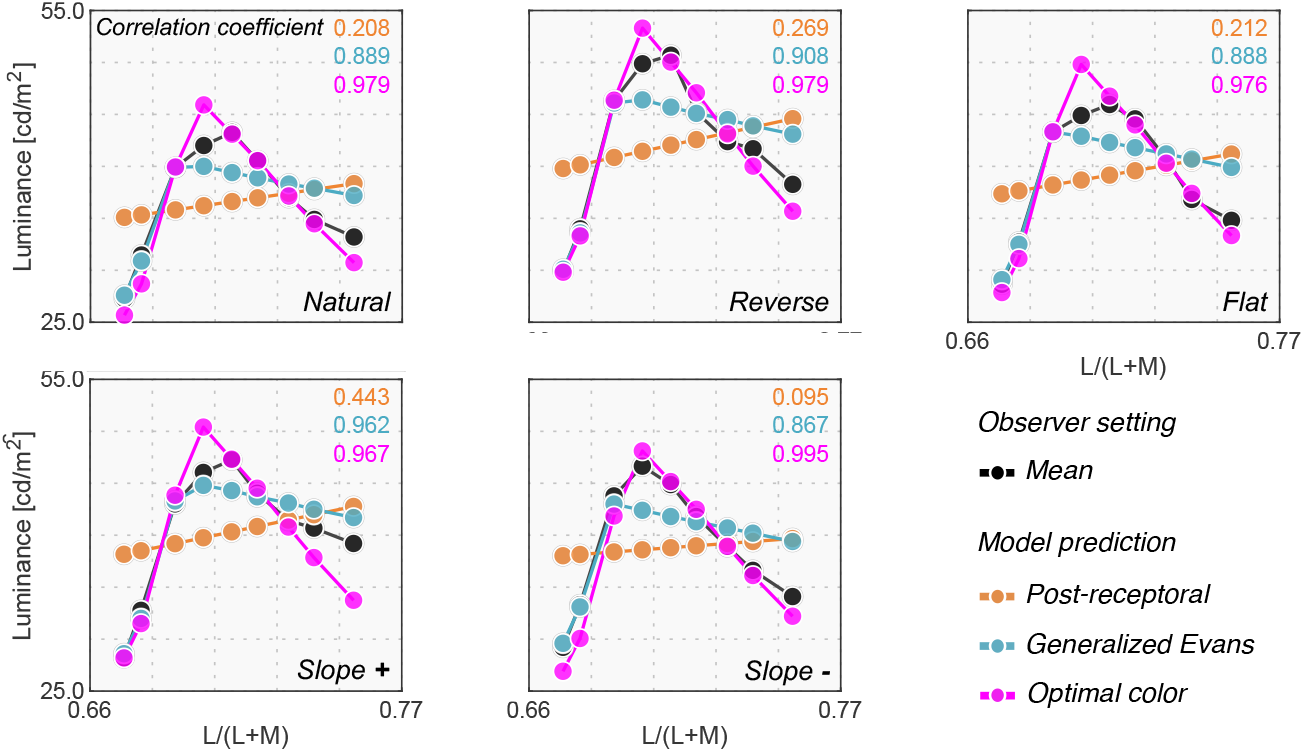
Predictions from the post-receptoral model, the generalized Evans’s model and the optimal color model in Experiment 1. Each model prediction was scaled to give the minimum root mean square error between model prediction and mean observer setting to compare their shapes more easily. The correlation coefficients between model prediction and mean observer settings are shown at the top right corner in each panel (in the order of post-receptoral model, generalized Evans’s model and optimal color model from top to bottom). We used a ground-truth illuminant (i.e., 6500K) to obtain the prediction from the optimal color model. [1.5-column figure]

**Figure A3:**
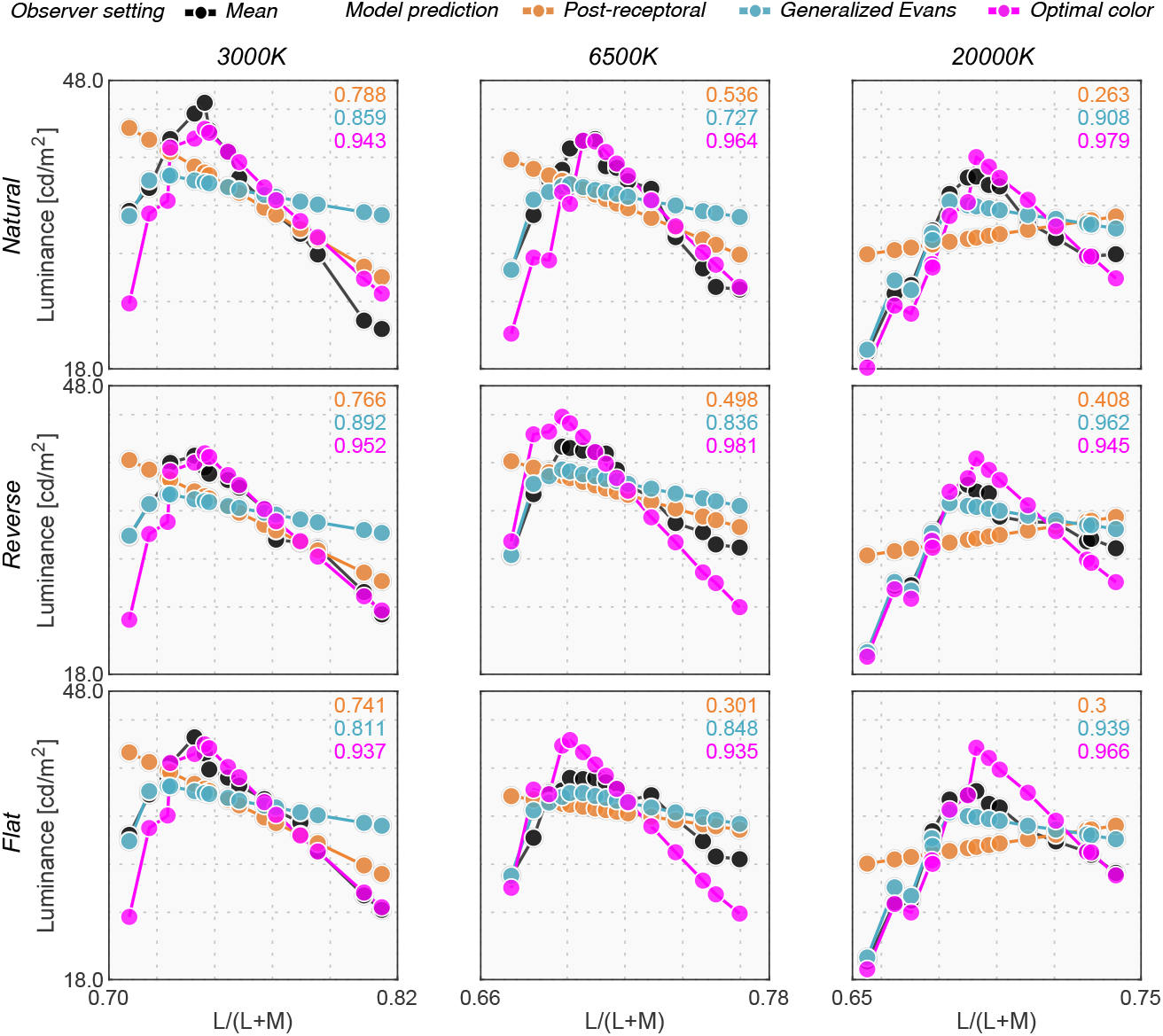
Predictions from the post-receptoral model, the generalized Evans’s model, and the optimal color model in Experiment 2. Each model prediction was scaled to give the minimum root mean square error between model prediction and mean observer setting. The correlation coefficients between model prediction and mean observer setting are shown at the right top corner in each panel. For the optimal color model, we used an estimated illuminant whose peak matched that of the observer settings (see Results section in Experiment 2 in the main text for more details). [1.5-column figure]

**Figure A4:**
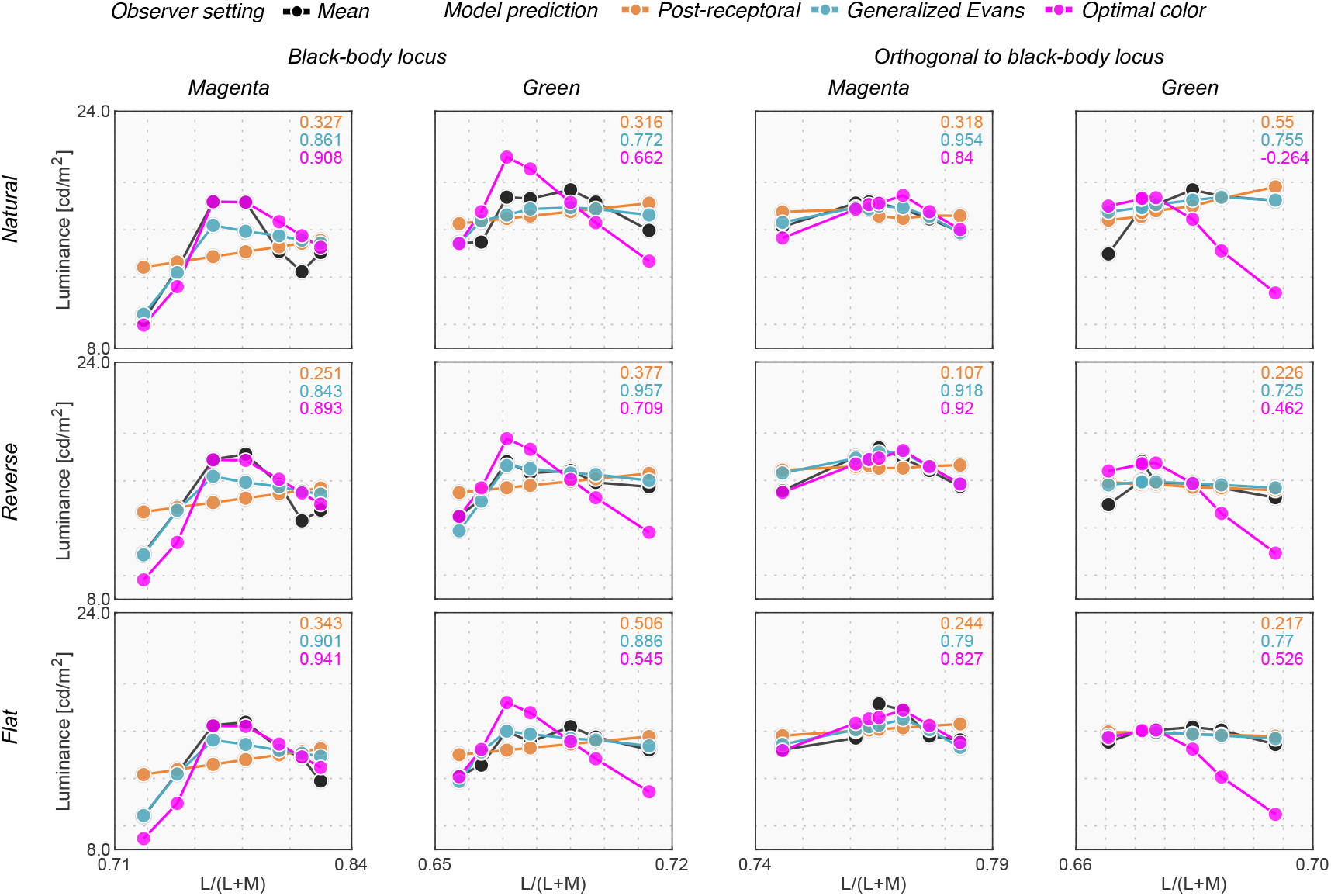
Predictions from the post-receptoral model, the generalized Evans’s model, and the optimal color model in Experiment 3. Each model prediction was scaled to give the minimum root mean square error between model prediction and mean observer setting. The correlation coefficient between model prediction and mean observer setting is shown at the top right corner in each panel. We used a ground-truth illuminant (i.e. magenta or green) for the optimal color model. [two-column figure]

